# Endosomal removal and disposal of dysfunctional, immunostimulatory mitochondrial DNA

**DOI:** 10.1101/2022.10.12.511955

**Authors:** Laura E. Newman, Nimesha Tadepalle, Sammy Weiser Novak, Cara R. Schiavon, Gladys R. Rojas, Joshua A. Chevez, Ian Lemersal, Michaela Medina, Sienna Rocha, Christina G. Towers, Danielle A. Grotjahn, Uri Manor, Gerald S. Shadel

## Abstract

Maternally inherited mitochondrial DNA (mtDNA) encodes essential subunits of the mitochondrial oxidative phosphorylation system, but is also a major damage-associated molecular pattern (DAMP) that engages innate immune sensors when released into the cytoplasm, outside of cells or into circulation^1^. This function of mtDNA contributes to antiviral resistance, but unfortunately also causes pathogenic inflammation in many disease contexts^2^. Cells experiencing mtDNA stress due to depletion of the mtDNA-packaging protein, Transcription Factor A, Mitochondrial (TFAM), or HSV-1 infection exhibit elongated mitochondria, mtDNA depletion, enlargement of nucleoids (mtDNA-protein complexes), and activation of cGAS/STING innate immune signaling via mtDNA released into the cytoplasm^3^. However, the relationships between altered mitochondrial dynamics and mtDNA-mediated activation of the cGAS-STING pathway remain unclear. Here, we show that entire enlarged nucleoids are released from mitochondria that remain bound to TFAM and colocalize with cGAS. These nucleoids arise at sites of mtDNA replication due to a block in mitochondrial fission at a stage when endoplasmic reticulum (ER) actin polymerization would normally commence, which we propose is a fission checkpoint to ensure that mtDNA has completed replication and is competent for segregation into daughter mitochondria. Released nucleoids also colocalize with the early endosomal marker RAB5 as well as the late endosomal marker RAB7 in TFAM-deficient cells and in response to mtDNA stress caused by the HSV-1 UL12.5 protein. Loss of RAB7 increases interferon stimulated gene (ISG) expression. Thus, we propose that defects in mtDNA replication and/or segregation enact a late mitochondrial fission checkpoint that, if persistent, leads to selective removal of dysfunctional nucleoids by a mitochondrial-endosomal pathway. Early steps in this pathway are prone to mtDNA release and cGAS-STING activation, but the immunostimulatory mtDNA is ultimately disposed of through a mechanism involving RAB7-containing late endosomes to prevent excessive innate immune signaling. This mtDNA quality control pathway might represent a therapeutic target to prevent mtDNA-mediated inflammation and associated pathology.

---

The vast majority of the mitochondrial proteome comprises proteins encoded by nuclear DNA, which includes all the enzymes and factors needed for replication of mtDNA and its packaging into nucleoids^4^. For example, TFAM is a dual HMG-box, mtDNA-binding protein essential for robust transcription initiation, transcription-primed mtDNA replication, nucleoid formation, and mtDNA segregation during mitochondrial fission^5–7^. We showed previously that haploinsufficency (*Tfam^+/−^*) or partial depletion of TFAM results in mtDNA depletion, enlarged nucleoids, elongated mitochondria (indicative of enhanced mitochondrial fusion and/or inhibited fission) and release of mtDNA into the cytoplasm that activates a cGAS-STING innate immune signaling pathway, characterized by upregulation of a unique subset of interferon-stimulated genes (ISGs)^3^. This response heightens antiviral immune responses in cells and mice and interestingly also enhances nuclear DNA repair capacity,^8^ showing that mtDNA release primes antiviral immunity and may also act as a cellular genotoxic stress sentinel that protects the nuclear genome^9^. To begin to unravel how altered mitochondrial and mtDNA dynamics cause mtDNA release and innate immune signaling, we first determined the nature of the cytoplasmic mtDNA in *Tfam^+/−^* MEFs. We performed live-cell imaging in *Tfam^+/−^* MEFs transfected with TOMM20-mCherry to mark the boundaries of the outer mitochondrial membrane (OMM) and co-stained with the DNA-intercalating dye Pico Green to label mtDNA. Time-lapse imaging of these cells revealed that a subset of the enlarged nucleoids is released from mitochondria in their entirety (Fig. 1A and Supplementary Video 1). To determine if these released nucleoids colocalize with cGAS, we repeated this experiment using fluorescently tagged cGAS (cGAS-mCherry) and also generated TFAM-deficient U2OS cells using CRISPR (Extended Data Fig. 1), to facilitate transfection and expression of multiple plasmids for imaging. These experiments reveal that enlarged nucleoids colocalize with cGAS-mCherry when they extend beyond the confines of the OMM (labeled by EBFP-Fis1[129-155]; Fig. 1B and Supplementary Video 2). Since Pico Green does not label mtDNA specifically, we confirmed these were in fact entire mtDNA nucleoids using fluorescence *in situ* hybridization (FISH) with previously characterized probes that target multiple regions of human mtDNA^10^ (Fig. 1C-F), performed in human primary lung fibroblasts (IMR-90) depleted of TFAM with siRNA. Furthermore, immunofluorescence performed after mtDNA FISH demonstrated that endogenous cGAS is localized with these now cytoplasmic, enlarged nucleoids (Fig. 1C-F), consistent with our time-lapse data in live cells using fluorescently-tagged cGAS in U2OS cells. And, although the TFAM signal was weak and often difficult to detect in IMR-90 cells transfected with TFAM siRNA, we consistently observed TFAM co-localization with released nucleoids and cGAS (Extended Data Fig. 2). Enlarged nucleoids also occur during infection by several viruses^3,11^, including HSV-1, and expression of the virus-encoded HSV-1 UL12.5 protein causes enlarged nucleoids before it eventually depletes mtDNA^3,12,13^. When expressed in U2OS cells, UL12.5 depleted mtDNA and reduced TFAM levels as we showed previously^3^, and also caused release of entire enlarged nucleoids that colocalize with cGAS and TFAM (Fig. 1G-J). Altogether, these results demonstrate that entire nucleoids are released under multiple mtDNA-stress conditions and that these remain bound to TFAM. Longer DNA molecules that are bound and extensively bent by the HMG-boxes of TFAM have been proposed to be optimal substrates for cGAS binding and activation^14^.

**Figure 1.**
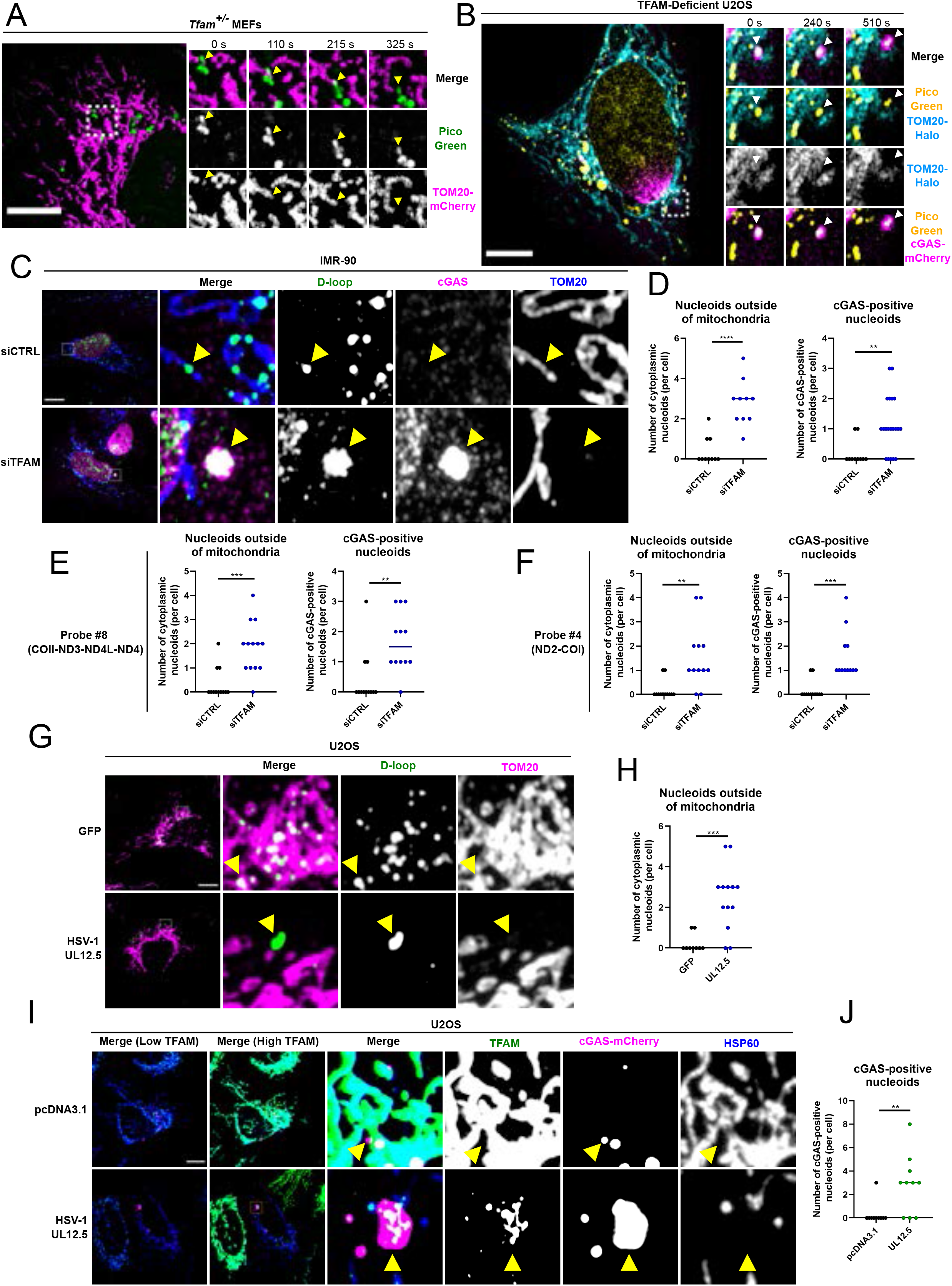
Mitochondrial DNA stress due to TFAM depletion or HSV-1 UL12.5 leads to release of entire enlarged nucleoids. **A)** Time-lapse airyscan imaging of Pico Green (DNA) and TOM20-mCherry (OMM) in a live, immortalized *Tfam^+/−^* MEF. Results were reproducible across two independent experiments. **B)** Time-lapse airyscan imaging of Pico Green (DNA), cGAS-mCherry, and TOM20-Halo (OMM) in a live TFAM-deficient U2OS cell. Results were reproducible across four independent experiments. **C)** Airyscan imaging of mtDNA FISH (D-loop probe), followed by immunofluorescence against cGAS and TOM20 in IMR-90 cells that were transfected with control or TFAM siRNAs. **(D-F)** Quantification of cytoplasmic nucleoids. The TOM20 channel was masked and subtracted from the mtDNA FISH channel to remove signals from nucleoids within mitochondria. The remaining cytoplasmic nucleoids per cell were counted (left) and scored for the presence of cGAS (right). This analysis was performed using mtDNA FISH probes to the D-loop region **(D)**, COII-ND4 (probe #8) **(E)**, or ND2-COI (probe #4) **(F).** N numbers are as follows: **(D)** (left) N=10 for both conditions, (right) siCTRL N=10, siTFAM N=20; **(E)** (left) siCTRL N =12, siTFAM N=13, (right) N=12 for both conditions; (F) (left and right) N=12 for both conditions. **G)** Airyscan imaging of mtDNA FISH (D-loop probe), followed by immunofluorescence against TOM20 in U2OS cells that were transfected with HSV-1 UL12.5 or GFP as a negative control. **H)** Quantification of cytoplasmic nucleoids present outside of mitochondria. N=9 for GFP, N=13 for UL12.5. **I)** Airyscan imaging of TFAM and HSP60 immunofluorescence in U2OS cells transfected with cGAS-mCherry and either pcDNA3.1 or HSV-1 UL12.5. For the cGAS-mCherry channel only, image display settings were not held constant between pcDNA3.1 and UL12.5, to allow for visualization of cGAS-mCherry structures and accounting for differences in the level of exogenous expression. **J)** Quantification of I, using TFAM as a marker for mtDNA and performed similarly to quantification in D. N=10 for both conditions. For all panels, scale bars = 10 μm, and for all plots lines represent mean.

We next probed the nature of the mtDNA and mitochondrial defects that are prompting the release of nucleoids from mitochondria. We first determined that the enlarged nucleoids in TFAM-depleted MEFs incorporate less 5-ethynyl-2’-deoxyuridine (EdU) than their smaller, more typical counterparts in the same cells (Fig. 2A,B). The diminished EdU signal in TFAM-depleted cells was not simply due to the lower overall mtDNA levels, as EdU intensity negatively correlated with nucleoid size (Extended Data Fig. 3). To determine if enlarged nucleoids localize to sites of mtDNA replication, we performed immunofluorescence against components of the mtDNA replisome, and observed that Twinkle, mtSSB, and POLG2 all colocalized with enlarged nucleoids (Fig. 2C). These results suggest that enlarged nucleoids comprise mtDNA molecules that are stalled after replication initiation or that have nearly completed a round of replication, but not yet segregated and still have the replication machinery associated. The ER licenses mtDNA replication at mitochondria/ER contact sites^15^. To determine whether the ER is present near enlarged nucleoids, we performed live cell imaging using Pico Green (DNA) in conjunction with markers of the ER (Ii33-mCherry) and mitochondria (mtBFP). Time-lapsed imaging shows that the ER wraps around those enlarged nucleoids that remain inside mitochondria in TFAM-deficient U2OS cells, and that these contacts are stable over at least a ten-minute imaging window (Fig. 2D), consistent with enlarged nucleoids arising at mtDNA replication sites and prolonged engagement of ER at these sites of mtDNA replication stress. Since mitochondrial fission is also coupled to mtDNA replication at mitochondria/ER contact sites^15^, we next asked if incomplete mtDNA replication prevents mitochondrial fission from proceeding, which could explain the mitochondrial elongation phenotype in TFAM-depleted cells. After ER association with mitochondria, actin polymerization on both mitochondria and ER are required for mitochondrial fission to proceed, with mitochondrial actin polymerization occurring first^16–22^. Using previously characterized organelle-specific actin probes^16^, we observed enrichment of mitochondrial-associated actin at the sites of enlarged nucleoids (Fig. 3A, quantified in 3C), but no obvious accumulation of ER-associated actin (Fig. 3B, quantified in 3D). These data indicate that TFAM-deficient cells are halted at an intermediate stage of mitochondrial fission, between the completion of mtDNA replication and initiation of ER actin polymerization. Consistent with this model, knockout of the ER-localized formin INF2, which polymerizes actin during mitochondrial fission^18,19^, caused enlarged nucleoids in U2OS cells (Fig. 3E,F), indicating that loss of ER-associated actin prevents mitochondrial fission and segregation of replicated nucleoids. Furthermore, loss of DRP1, which mediates the final steps of mitochondrial fission^23–25^, not only causes enlarged, “clustered” nucleoids^26^ (Fig. 3G,H), but also release of nucleoids that colocalize with cGAS (Fig. 3J,K) and enhanced expression of ISGs (Fig. 3I) in wild type cells, robustly phenocopying the loss of TFAM. Therefore, loss of mitochondrial fission prevents segregation of nucleoids, suggesting close coordination of mtDNA replication and fission, and inhibition of either process stalling the other. In addition, both mitochondrial perturbations can cause clustering and cytoplasmic release of nucleoids and cGAS-STING activation.

**Figure 2.**
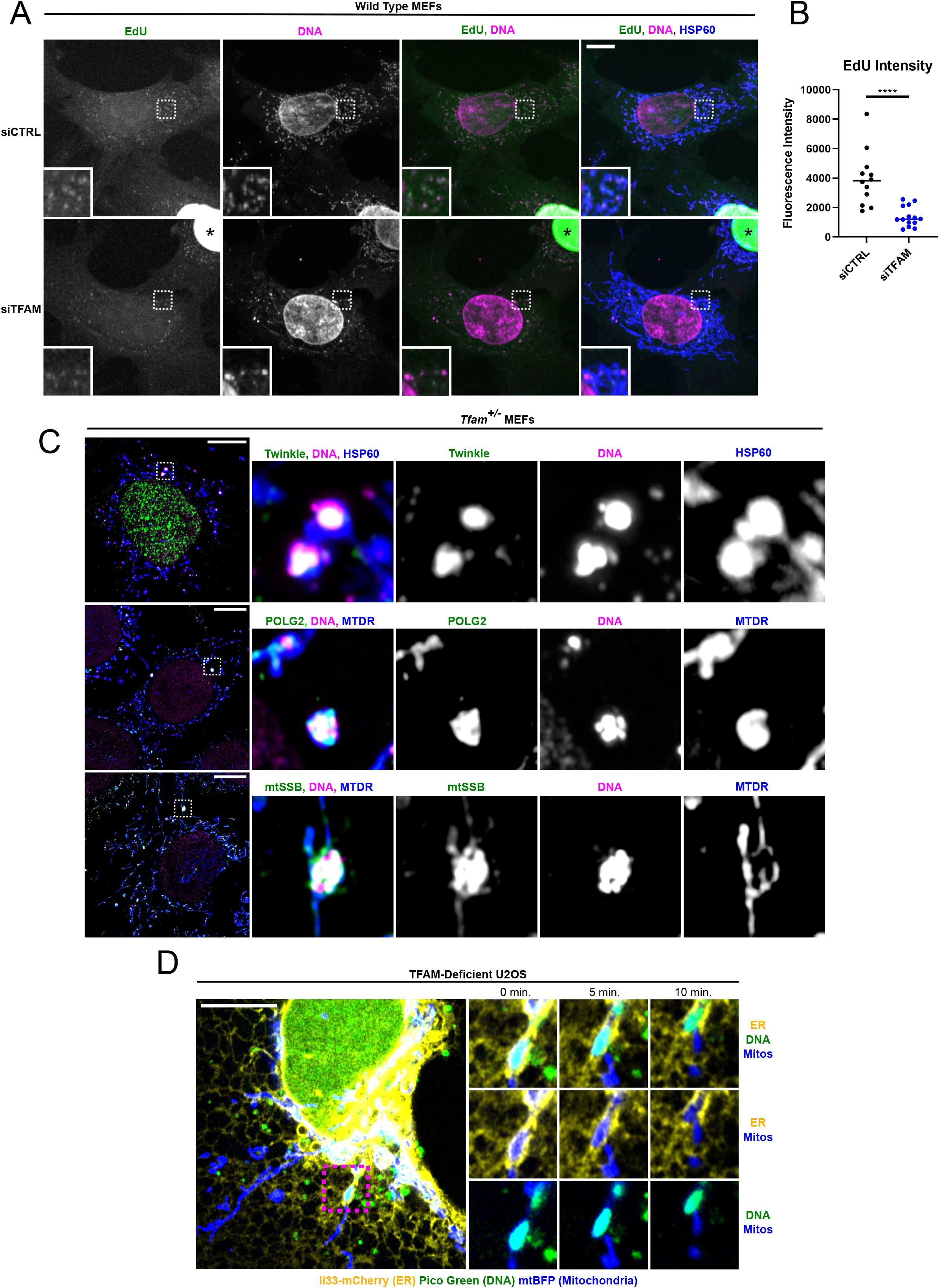
Incomplete mtDNA replication causes nucleoid clustering and prolonged mitochondria/ER contacts. **A)** Pulse EdU labeling (4 hours) of immortalized wild-type MEFs that were previously transfected with control or TFAM siRNAs for 96 hours. Asterisk (*) denotes EdU incorporation within the nuclei of some cells in the population, shown as an internal positive control. Confocal images of Alexa488-EdU, DNA, and HSP60 are shown. **B)** Quantification of **(A)**. A mask of nucleoids was created by thresholding non-nuclear DNA, and EdU fluorescence intensity was measured within the mask. N=12 for siCTRL, N=14 for siTFAM. **C)** Airyscan imaging of immunofluorescence against Twinkle, POLG2, or mtSSB alongside DNA in immortalized *Tfam^+/−^* MEFs. Mitochondria were labeled using either HSP60 or Mitotracker deep red (MTDR). Results were reproducible across two independent experiments. **D)** Airyscan imaging of a live TFAM-deficient U2OS cell (clone #2) of Ii33-mCherry (ER), Pico Green (DNA), and mtBFP (mitochondria). Time-lapse imaging demonstrates that mitochondria/ER contacts around enlarged nucleoids are stable over a ten-minute imaging window. Results were reproducible across three independent experiments. For all panels, scale bars = 10 μm, and for all plots line represents mean.

**Figure 3.**
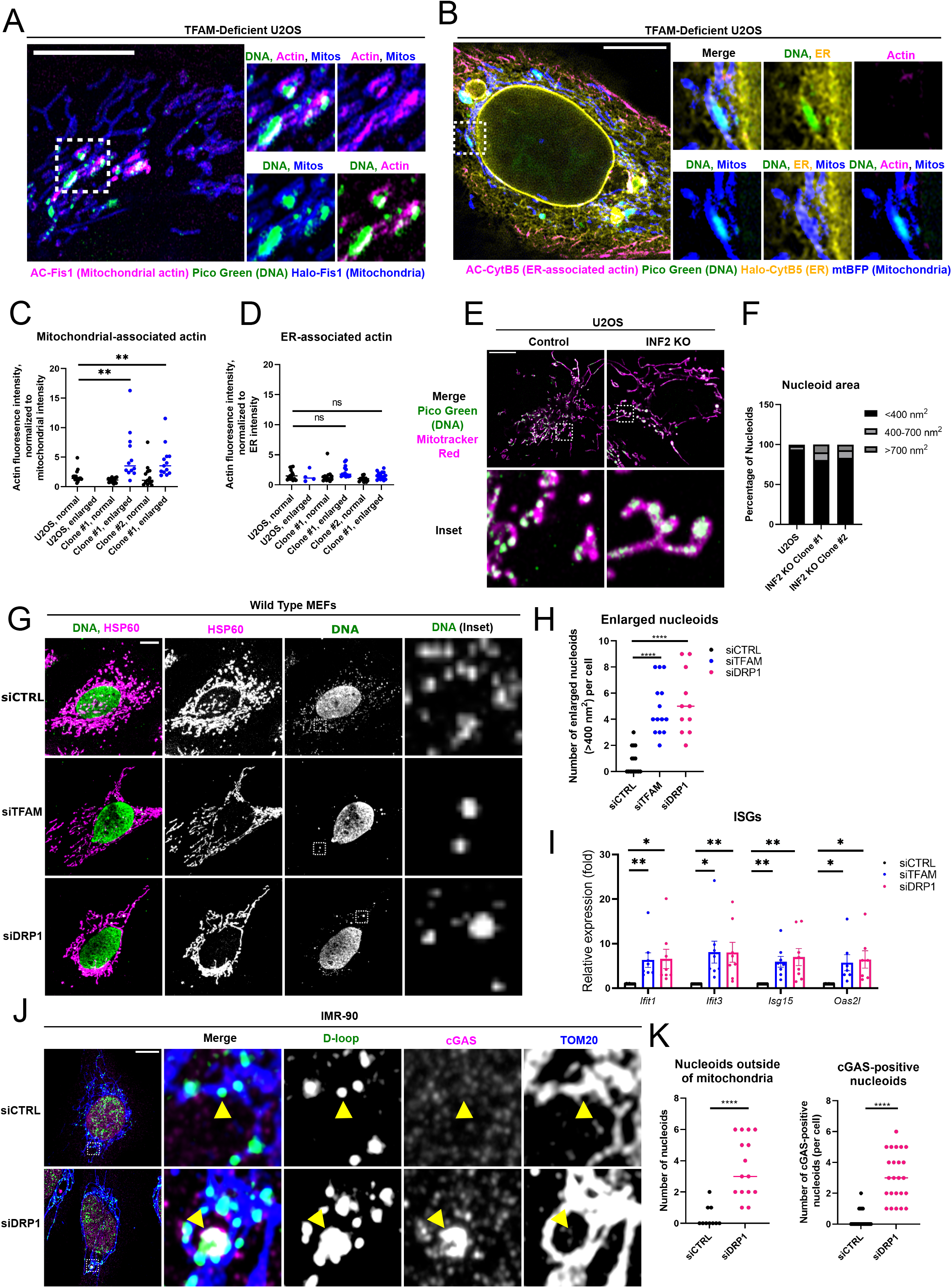
A mitochondrial fission checkpoint coupled to mtDNA replication. **A)** Airyscan imaging of a live TFAM-deficient U2OS cell (clone #1) expressing a mitochondrially targeted actin nanobody (AC-mCherry-Fis1) to visualize actin accumulation on mitochondria alongside Pico Green (DNA) and the mitochondrial outer membrane (OMM) (Halo-Fis1). **B)** Airyscan imaging of a live TFAM-deficient U2OS cell (clone #1) expressing an ER targeted actin nanobody (AC-mCherry-CytB5) to visualize actin accumulation on the ER alongside Pico Green (DNA), the mitochondrial matrix (mtBFP), and the ER (Halo-CytB5). **(C,D)** Quantification of F and G. Line scans were drawn through normal or enlarged nucleoids within mitochondria, and the peak actin fluorescence intensity was measured and normalized to mitochondria or ER fluorescence intensity. For all conditions, N=15 nucleoids across 5 cells. **E)** Airyscan imaging of Pico Green (DNA) and Mitotracker red (mitochondria) in live U2OS cells. **F)** Quantification of **(E).** N=17 cells for each condition. **G)** Confocal imaging of primary wild type MEFs transfected with siRNAs against TFAM or DRP1. Immunofluorescence against DNA and HSP60 is shown. **H)** Quantification of nucleoids in panel C. The number of nucleoids larger than 0.4 um^2^ per cell was quantified by thresholding the non-nuclear DNA signal and measuring particle size using ImageJ. N=16 for siCTRL, N=14 for siTFAM, and N=11 for siDRP1. **I)** qRT-PCR of primary wild type MEFs transfected with TFAM or DRP1 siRNAs. Eight biological replicates (dots) are shown as mean ± standard error of the mean. **J)** Airyscan imaging in a DRP1-depleted IMR-90 cell of mtDNA FISH D-loop probe, followed by immunofluorescence against cGAS and TOM20. **K)** Quantification of cytoplasmic nucleoids (N=10 for siCTRL, N=15 for siDRP1) and cGAS-positive nucleoids (N=22 for siCTRL, N=24 for siDRP1) as in Figure 1D. All scale bars = 10 μm, and for all plots line represents mean.

Based on the results above, we propose that the enlarged nucleoids in TFAM-depleted cells arise from a failure to complete mtDNA replication and/or segregation which halts mitochondrial fission as part of a late checkpoint to ensure inheritance of mtDNA molecules in each daughter mitochondria during mitochondrial division. We next hypothesized that prolonged engagement of this mitochondrial fission checkpoint enacts an adaptive response to remove these defective nucleoids that is prone to cytoplasmic mtDNA release and cGAS-STING innate immune activation. One mechanism for removal/release of entire mtDNA nucleoids that also activates cGAS-STING signaling is through large BAK/BAX pores under conditions of caspase-inhibited apoptosis^27,28^. However, this does not appear to be occurring in *Tfam^+/−^* MEFs, as we were unable to detect the presence of BAX pores in *Tfam^+/−^* MEFs (Extended Data Fig. 4A,B), did not observe any obvious inner membrane herniation around enlarged nucleoids (Extended Data Fig. 4C) and still observed cytoplasmic nucleoids in TFAM-depleted *Bak^−/−^Bax^−/−^* MEFs (Extended Data Fig. 4D,E). Another mechanism involves release of mtDNA fragments through pores comprised of VDAC1 or VDAC3 oligomers^29^. Though this mechanism does not explain release of entire nucleoids, we nonetheless decided to test for the role of VDAC pores in release of enlarged nucleoids, in case shorter mtDNA fragments were also being released and driving cGAS/STING signaling in *Tfam^+/−^* MEFs. Depletion of VDAC1 or VDAC3 had no effect on ISGs in TFAM-deficient cells (Extended Data Fig. 5A,B), and we still documented cytoplasmic nucleoids in IMR-90 cells in which both TFAM and VDAC1 were depleted (Extended Data Fig. 5C,D). Thus, we next endeavored to capture mtDNA release events by correlative light and electron microscopy (CLEM) to better understand the nucleoid release mechanism in TFAM-depleted cells. Live TFAM-deficient U2OS cells labeled with Pico Green, cGAS-mCherry, TOM20-Halo, and mTurquoise-LC3 (for fiducial information) were imaged using airyscan microscopy until an apparent mtDNA-release event in progress was observed, at which point the cell was immediately fixed on the microscope stage. The same cell was then located, serially sectioned, and imaged using scanning electron microscopy (SEM), after which the data sets were aligned (Extended Data Fig. 6A). We were able to locate a couple of structures that were positive for Pico Green and cGAS-mCherry, but negative for TOM20-Halo, including one structure apparently budding from a TOM20-positive mitochondrion (Extended Data Fig. 6B-E), indicating enlarged nucleoids can be extruded from mitochondria within a membrane-bound compartment. Unlike previously described IMM herniations, these compartments did not appear to be extruding through a break in the outer mitochondrial membrane. Mitochondrial-derived vesicles (MDVs) can bud from mitochondria and carry mitochondrial cargo to the lysosome^30^. However, Pico Green and cGAS-mCherry positive structures are much larger than IMM-derived MDVs, which are ~100 nm in diameter^30,31^. Therefore, the nucleoid-containing structures in TFAM-depleted cells appear to represent membrane-bound compartments other than MDVs.

A clue to the nature of these mtDNA-containing vesicular compartments came from parallel, cryo-electron tomography experiments in *Tfam^+/−^* MEFs, in which we consistently observed multi-membrane structures that resembled multivesicular bodies (Fig. 4A,B), which is a type of late endosome^32^. An example cryo-EM tomogram slice is shown (Fig. 4A), and this data was reconstructed and segmented data for easier visualization (Extended Data Fig. 7). We observed multivesicular bodies in CLEM experiments as well and noted that cGAS-mCherry fluorescence signal overlapped with multivesicular bodies in *Tfam^+/−^* MEFs (Extended Data Fig. 6G-I). Therefore, we hypothesized that *Tfam^+/−^* MEFs are experiencing increased flux through the endosomal system, and that this pathway may be involved in selective removal and degradation of the aberrant nucleoids. To test this, we repeated mtDNA FISH experiments in combination with immunofluorescence against the late endosomal marker RAB7, and found that RAB7 overlapped with cGAS-positive nucleoids located outside of mitochondria (Fig. 4C,D). We also found that RAB7A-GFP overlapped with cytoplasmic TFAM and cGAS-mCherry in U2OS cells expressing the HSV-1 UL12.5 protein (Fig. 4E,F), demonstrating that this pathway is not unique to TFAM depletion per se. With regard to innate immune signaling, depletion of RAB7 increased ISGs above the levels normally present in *Tfam^+/−^* MEFs (Fig. 4H), and the fold increase in ISGs with RAB7 siRNA was higher in *Tfam^+/−^* cells than in wild-type MEFs, suggesting that loss of RAB7 activity prevents maturation and disposal of endosomes containing cGAS-bound mtDNA, thereby prolonging mtDNA-cGAS-STING signaling. However, the question of how nucleoids are extracted and initiate cGAS-STING signaling in the first place remained. One possibility is that mitochondria containing enlarged nucleoids could be targeted for degradation by mitophagy, which could explain why released mtDNA colocalizes with RAB7, as RAB7 also marks lysosomes, which fuse to autophagosomes during the final steps of autophagy. However, mitophagy flux in cells expressing HSV-1 UL12.5 was similar to control cells (Extended Data Fig. 8). Therefore, we sought another explanation for how mtDNA arrives in a RAB7 compartment. Early endosomes are regulated by RAB5 and mature into late endosomes in a RAB7-dependent manner^33^. We therefore hypothesized that this pathway might also initiate at mitochondrial membranes and mediate extraction of the offending nucleoids into endosomes (Fig. 4G). In support of this idea, RAB5 is recruited to the mitochondrial OMM in several contexts, including mitophagy and oxidative stress^34–36^. We observed enlarged nucleoids colocalized with mCherry-RAB5 outside of mitochondria (Fig. 4I,J), indicating that enlarged nucleoids traffic within early endosomes, explaining their arrival at late endosomes. However, it appears that the amount of cGAS-STING signaling is kept largely in check by movement of the immunostimulatory nucleoids into late endosomes for ultimate disposal.

**Figure 4.**
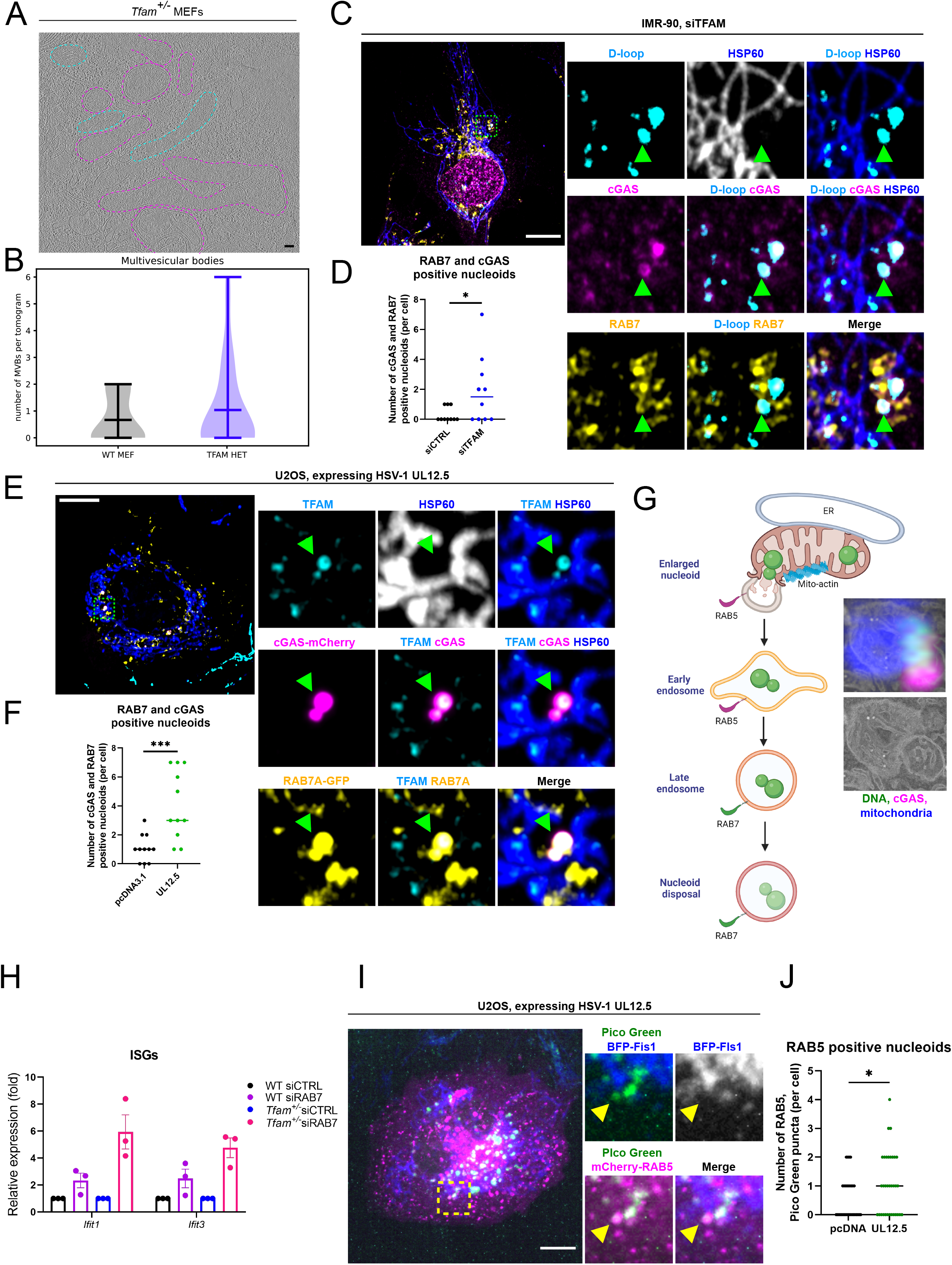
Dysfunctional nucleoids are removed and degraded by the endolysosomal pathway to prevent excessive innate immune signaling. **A)** A representative tomogram slice of a *Tfam^+/−^* MEF cell imaged by cellular cryo-electron tomography. Multivesicular bodies are highlighted by dashed pink lines, and mitochondria were highlighted by dashed cyan lines. Scale bar = 100 nm. **B)** The presence of multivesicular bodies was scored as described under methods. **C)** Airyscan imaging of mtDNA FISH D-loop probe, followed by immunofluorescence against RAB7, cGAS, and HSP60 in TFAM-depleted IMR-90 cells. **D)** The number of cytoplasmic nucleoids per cell that were positive for both cGAS and RAB7 was scored. N=10 for each condition. **E)** Airyscan imaging of TFAM and HSP60 immunofluorescence in U2OS cells expressing RAB7A-GFP, cGAS-mCherry, and HSV-1 UL12.5. **F)** The number of TFAM-marked nucleoids per cell that were positive for both cGAS and RAB7 was scored. N=11 for both conditions. **G)** Schematic summarizing endolysosomal extraction and degradation of enlarged nucleoids from mitochondria, where RAB5 drives the formation of early endosomes, which then mature and dispose of nucleoids in a RAB7-dependent manner. CLEM data of mtDNA release from Extended Data Fig. 6 is shown for comparison. **H)** qRT-PCR of primary wild type or *Tfam^+/−^* MEFs transfected with control or RAB7 siRNA, normalized to beta actin. Dots represent biological replicates (N=3). For both wild type and *Tfam^+/−^* MEFs, ISGs were normalized to siCTRL, in order to compare fold increase with each siRNA. **I**) Spinning disk imaging of Pico Green (DNA), BFP-Fis1 (mitochondria), and mCherry-RAB5B (early endosomes) in live cells expressing HSV-1 UL12.5. J) Quantification of released nucleoids positive for mCherry-RAB5B, as in I (N=31 for pcDNA, N=34 for UL12.5). Data are reported as mean ± standard error of the mean. All scale bars = 10 μm, and for all plots line represents mean.

Our results define multiple connections between mtDNA replicative stress, mitochondrial fission and innate immune activation. While it is known that the ER couples mtDNA replication to mitochondrial fission to maintain distribution of mtDNA within the mitochondrial network^15^, we show here that disruption of this process leads to an adaptive response to selectively remove non-functional nucleoids by a mitochondrial-endosomal pathway that is prone to mtDNA release and involves RAB5, RAB7 and likely other components involved in early endosome formation on the mitochondrial surface (Extended Data Fig. 9). In the initial stages of this response, cGAS localizes to the released nucleoids and innate immune signaling commences, but excessive innate immune signaling is prevented by movement of the immunostimulatory nucleoids into late endosomes for ultimate disposal. Interestingly, this response can be initiated by stalled mtDNA replication that leads to inhibited fission, or by inhibited DRP1/ER-mediated fission that stalls mtDNA replication/segregation and leads to nucleoid clustering and release. That is, there is bidirectional signaling occurring between ER and mtDNA replication status that provides a mtDNA checkpoint for mitochondrial fission.

There is growing evidence for additional pathways beyond mitophagy to maintain mitochondrial quality control, including MDVs^31^ as well as piecemeal autophagy of mitochondrial components such as nucleoids^37^ and MICOS complexes^38^. Importantly, it was recently demonstrated that mitophagy suppresses mtDNA-driven inflammation^39^. Therefore, how these additional mitochondrial quality control pathways interface with innate immune pathways is worthy of additional study. Very recent evidence indicates that the endolysosomal pathway also mediates mitochondrial quality control, either by enhancing mitophagy^34,40^, or by mediating mitochondrial clearance in the absence of autophagy^41^. We demonstrate a role for the endosomal system in the disposal of nucleoids that fail to finish replication and/or segregation, elucidating another pathway of selective disposal of nucleoids, separate from mitophagy^37^. Finally, we propose that endolysosomal extraction of nucleoids represents a third mtDNA release pathway separate from that mediated by BAK/BAX^27,28^ or VDAC pores^29^. Therefore, it will be important to fully characterize which mtDNA-release mechanisms operate under different physiological and pathological contexts. Importantly, endolysosomal extraction of enlarged nucleoids also occurs when HSV-1 UL12.5 is expressed, indicating that the endosomal mtDNA-release pathway is likely relevant during HSV-1 viral infection and perhaps other infections that have been reported to cause mtDNA release^3,11,42–44^. Finally, our results point to mtDNA replication, mitochondrial fission, and the endolysosomal pathway as potential targets to prevent mtDNA release or augment disposal of immunostimulatory mtDNA to prevent inflammation pathology associated with multiple diseases and aging.

## Experimental Procedures

### Animal strains and cell lines

*Tfam^+/−^* mice were originally derived from *Tfam^flox^* mice obtained from N. Chandel (Northwestern University) and generated as described previously^1,2^. All animal husbandry and procedures were Institutional Animal Care and Use Committee (IACUC) approved by the animal care and use committees at Yale University or the Salk Institute for Biological Studies. IMR-90 human fetal lung fibroblasts and *Bak^−/−^Bax^−/−^* MEFs were purchased from ATCC. TFAM-deficient and INF2 knockout U2OS cell lines were derived from a CRISPR-knockout pool of U2OS cells purchased from Synthego by using limiting dilution to obtain single cell clones. Editing of TFAM was confirmed by sequencing. The parental U2OS line from the ATCC was supplied alongside the CRISPR pooled cells and used in all experiments. All experiments using CRISPR-mediated KO were performed using two independent clones in order to guard against clonal as well as off-target CRISPR differences. For TFAM-deficient cells, clone #1 (clone number 5G8) and clone #2 (clone number 1C11) were used in all experiments. For INF2 KO cells, clone #1 was compared against clone #2, which was a gift from Henry Higgs^3^.

### Cell culture

Wild type and *Tfam^+/−^* MEFs were generated from mouse embryos isolated between embryonic day 12.5 and 14.5 and cultured in DMEM (Corning), supplemented with 10% fetal bovine serum (FBS) (Gibco, Thermo Fisher). All experiments were performed in primary MEFs passaged five or fewer times. IMR-90 cells were obtained from the ATCC and were cultured in EMEM supplemented with 10% FBS and were not used for experiments past 30 population doublings. Immortalized cell lines were not used for experiments beyond passage 30. TFAM-deficient and INF2 KO U2OS clones as well as parental U2OS cells were cultured in DMEM supplemented with 10% FBS. To generate stable TFAM shRNA cell strains, LentiCRISPRv2 vector with predesigned shRNA (Sigma mission shRNA TRCN0000312779, target sequence TGTCAAACTAGAACGGATAAA) was transfected into Lenti-x 293T cells with 2:1 (psPAX2/pMD2G) plasmids using Lipofectamine 2000 (Thermo Fisher # 12566014) at a ratio of 1:1 (Lipofectamine 2000/nucleic acid (μg)) to produce lentivirus. Lentivirus-containing media were collected 48 h post-transfection and filtered using a 0.45-μm membrane. *Bak^−/−^Bax^−/−^* (ATCC) or wild type immortalized MEFs were infected with lentivirus for 24 h and then selected for antibiotic resistance in 20 μg ml^-1^ blasticidin for 5 days. Fibronectin coating was achieved by incubating dishes with fibronectin (10 μg ml^-1^) diluted from stock solution (Sigma #F1141). For live cell imaging experiments, cells were stained with Pico Green (1:500, Thermo Fisher #P11495) and 100 nM of Mitotracker red or deep red (Thermo Fisher #M7512, M22426) (when indicated) for 30 min., washed 3x with PBS, and imaged in phenol-red free DMEM (Thermo Fisher # 21-063-029) plus prolong antifade (Thermo Fisher #P36975) along with Janelia Fluor 635 (250-500 nM) (if a Halo-tagged construct was expressed). For all experiments, cells were targeted for a final density of 50-70%, and variations in cell density between experimental conditions was minimized. All cells were screened monthly for mycoplasma using a kit. Additional mycoplasma screening was regularly performed by staining cells with Hoescht 33342 during imaging experiments, whenever possible.

### Transfection

Transfection of siRNAs (Extended data Table 2, purchased from IDT) into MEFs and IMR-90 at 50-70% density was achieved using Lipofectamine RNAiMax (Invitrogen #1847641) according to manufacturer’s instructions. All siRNAs were transfected at a final concentration of 25 nM. Following 24 hours of transfection, cells were passaged (typically 1:8 or 1:10) into new dishes or onto coverslips and harvested 96 hours from the start of transfection. For IMR-90 cells, both the new plates and coverslips were coated with freshly prepared fibronectin solution prior to passaging immediately after transfection. Transfection of plasmid DNA was performed using Lipofectamine 2000 (Thermo Fisher # 12566014) at a ratio of 4:1 (Lipofectamine 2000/nucleic acid (μg)), according to manufacturer’s instructions. Confluent U2OS cells were transfected with 2 μg in one well of a 6 well plate. When transfecting multiple plasmids, the amount of plasmid transfected was evenly split to reach a final amount of 2 μg for all plasmids, with the exception of fluorescent proteins with emission in the blue range, for which the amount of plasmid DNA was doubled relative to the other plasmids. Following 4 hours of transfection in growth media, cells were trypsinized and plated onto coverslips (1:2 split) or Nunc 8 well chamber slides (Thermo Fisher # 155409PK) (20-40k cells per well).

### Plasmids

The following plasmids were purchased from Addgene: TOMM20-mCherry (#55146), TOM20-Halo (#111135), mtBFP (#55248), AIF-mCherry (#67530), mTurquoise-LC3 (#55579), mCherry-RAB5B (#49201), psPAX2 (#12260), pMD2.G (#12259). AC-mito and ACER fused to mCherry were previously described^4^. Human cGAS fused to EGFP in pcDNA3.1(+) was purchased from Genescript, and mCherry was subcloned in place of EGFP. To generate EBFP-Fis1 and Halo-Fis1, EBFP or Halo was subcloned into the mCherry-Fis1 construct that was previously described (mCherry-mito^4^). HSV-1 UL12.5 was purchased from Addgene (#70109) and the EGFP was removed by subcloning. Ii33-mCherry was a generous gift from P. Satpute (National Institutes of Health, Bethesda, MD), and RAB7A-GFP^5^ was a generous gift from Chengbiao Wu (UCSD).

### Immunofluorescence and Fluorescence in situ hybridization (FISH)

Cells grown on fibronectin-coated coverslips (Fisher Scientific #12-545-81) were fixed at 37°C using a prewarmed (37°C) solution of 4% paraformaldehyde in PHEM buffer (60 mM PIPES, 25 mM HEPES, 10 mM EGTA, 4 mM MgSO_4_, pH 6.8) for 15 minutes, and then permeabilized with 0.1% (v/v) Triton X-100 in PBS for 10 minutes at room temperature. FISH was then performed as previously described^6^. Probes #4 and #8 were prepared as previously described, and the mREP probe conjugated to Atto550 or Atto633 was purchased from Integrated DNA Technologies^6^. For immunofluorescence, coverslips were blocked with filtered PBS containing 1% (w/v) at room temperature for at least an hour, following either permeabilization or FISH. Incubation with primary antibodies was carried out in PBS containing 1% (w/v) BSA at 4°C overnight, followed by 4 x 5 minute washes in PBS. Secondary antibodies (1:500) were incubated in PBS containing 1% BSA for 1 hour at room temperature. Secondary antibody was removed by 4 x 5 minute washes in PBS. Coverslips were then mounted onto slides using Prolong Glass. The following adjustments were made for particular experiments. If the anti-DNA primary antibody was used, washes following primary incubation were instead performed using 5 mM EDTA in PBS, as we found that it effectively removed background for that particular antibody. If FISH was performed prior to immunofluorescence, the concentration of primary antibodies was doubled, to account for loss of signal. When four color imaging was performed, HSP60 was imaged in the blue channel, and the concentration of both primary (HSP60) and secondary antibody (Alexa 405) was doubled. To visualize Twinkle, antigen retrieval was performed prior to permeabilization by heating coverslips to 95°C in antigen retrieval buffer (100 mM Tris, 5% [w/v] urea, pH 9.5) for 10 minutes, followed by 4 x 5 minute washes in PBS. Immunofluorescence was then performed using freshly purchased anti-Twinkle antibody.

The following antibodies were obtained commercially and used at the indicated dilutions for immunofluorescence: mouse monoclonal TOM20 (BD Biosciences #612278, 1:1000), TOMM20 (Abcam #ab186734, 1:200), cGAS (CST #15102, 1:100), TFAM (Proteintech #22586-1-AP, 1:500), TFAM (Abcam #ab119684, 1:1000), DNA (Millipore #CBL186, 1:200), HSP60 (CST #12165, 1:1000), HSP60 (EnCor Biotechnology #CPCA-HSP60, 1:1000), Twinkle (Abcam #ab83329, 1:100), POLG2 (Proteintech #10997-2-AP, 1:100), mtSSB (Proteintech #12212-1-AP, 1:100), RAB7 (CST #9367, 1:100), Activated BAX (Santa Cruz Biotechnology #sc-23959, 1:100). Secondary antibodies conjugated to Alexa 405, 488, 568, or 647 were purchased from Thermo Fisher, with the exception of anti-chicken Alexa 405 and 647, which were from Abcam.

### EdU incorporation

EdU experiments were performed using the Alexa 488 Click-It EdU kit (Thermo Fisher #C10637), with the following modifications to the manufacturer protocol. Cells on coverslips were incubated with EdU (50 μM) for 4 hours prior to fixation. After fixation, two sequential, hour long click reactions were performed, using a freshly made solution for each reaction. Coverslips were then washed 3x with 3% BSA in PBS, after which point immunofluorescence was performed as described above.

### Microscopy

Fixed cells were imaged with a Plan-Apochromat ×63/1.4 NA oil objective on an upright Zeiss 880 LSM Airyscan confocal microscope. For all fixed cell imaging, images shown are maximum intensity z projections, unless otherwise noted. Live cells were imaged with a Plan-Apochromat ×63/1.4 NA oil objective on an inverted Zeiss 880 LSM Airyscan confocal microscope with the environmental control system supplying 37 °C, 5% CO_2_ and humidity. For time-lapse imaging, the zoom factor was set to 3-4× to increase the frame rate. In all cases, the maximum pixel-dwell time (~0.684 μs per pixel) and 2× Nyquist optimal pixel size (~40 nm per pixel) was used. For all live cell images, single planes are shown. Airyscan images are denoted as “airyscan images” in the figure legends. For some experiments, imaging of fixed cells was performed using an HCX Plan-Apochromat ×100/1.4 NA CS oil objective on an inverted Leica SP5 microscope, these images are denoted as “confocal images” in the figure legends. For some experiments, imaging was performed using a Plan-Apochromat ×63/1.4 NA oil objective on an inverted Zeiss spinning disk microscope, these images are denoted as “spinning disk images” in the figure legends. For all experiments, 405, 488, 561, and 633 nm laser lines were used.

### Correlative live cell imaging and electron microscopy

TFAM-deficient U2OS cells were transfected with cGAS-mCherry, TOM20-Halo, and mTurquoise-LC3 and plated onto gridded coverslips in 35 mm dishes (Cellvis #D35-14-1.5GI). Cells were stained with PicoGreen and JF635. The PicoGreen, cGAS-mCherry, and TOM20-Halo channels were imaged live until an apparent mtDNA release event occurred, which defined ROIs for EM imaging. Samples were then fixed while imaging on the microscope by adding dropwise an equal volume of 2X EM-fixative (5% glutaraldehyde, 4% paraformaldehyde in 0.1M cacodylate buffer with 3mM CaCl_2_) warmed to 37°C to the live cell media already in the dish. After fixation, oversampled z stacks (100 nm step size) were collected of all four channels. Additional images of the surrounding cells and the grid were also acquired. Samples were immediately removed from the microscope, fixative-media solution was discarded, and cells were washed once and left in fresh ice cold 1X fixative for at least 60 minutes and rinsed three times with ice cold 0.1M cacodylate buffer with 3mM CaCl_2_. Coverslips were post-fixed and stained with 1.5% reduced osmium for 35 minutes, rinsed five times with MilliQ water, and stained again with 1% aqueous uranyl acetate for 1 hour at room temperature, before serial dehydration with graded solutions of ice-cold ethanol in water. Samples were then fully dehydrated in two 10 minute rinses of anhydrous ethanol at room temperature before infiltration with Durcupan resin. After 3:1, 1:1, 1:3 ethanol-resin two-hour infiltration steps, samples were infiltrated with pure resin for two hours before another change of fresh pure resin and left overnight at room temperature. In the morning, a final change of fresh resin was performed, taking care to let the viscous resin fully drain by inverting the coverslip before adding the final aliquot of fresh resin from the edge of the dish. Resin was added to fill the dish up to 2 mm and infiltrated samples were polymerized for 48 hours at 65°C. After polymerization, coverslips were dissolved by immersion in concentrated hydrofluoric acid. Correlative light-electron microscopy (CLEM) was achieved using laser branded fiducials in a thinly embedded sample, as previously described^7^. Briefly, regions of interest (ROIs) were identified by their grid positions and dark osmium staining. ROIs were marked using the cutting laser of a Zeiss PALM laser cutting microscope to provide orientation and fiducial guides for further trimming and ultramicrotomy. Following laser cutting, an additional protective layer of resin was applied. ROIs were carefully identified under a dissecting microscope and excised from the coverslip using a jeweler saw and a scalpel, with careful thought given to the future blockface orientation. The small (1×2×2mm) sample block was glued to a blank resin block such that the ROIs were orthogonal to the cutting plane and approximately 100 μm from the cutting surface. Laser marks on the block face helped to identify the location of the ROI using the ultramicrotome optics. The block was carefully trimmed using a 90° diamond trimming knife (Diatome) so that the block width bounded the fiducial laser marks, and the two sides of the block face were perfectly parallel for serial sectioning when turned 90°. The ROI was approached using the trimming knife to provide a perfectly smooth block face. When the fiducials marking the location of ROI became challenging to visualize (about 5-15μm from the blockface), a diamond knife with a water boat appropriate for serial sectioning (Histo 45°, Diatome) was installed on the ultramicrotome and 70 nm serial sections were collected on a series of plasma cleaned custom-diced silicon chips (University Wafer, Boston, MA) immersed in the knife boat. The chips were mounted on aluminum stubs using double-sided carbon sticky tape and loaded into a Zeiss Sigma VP scanning electron microscope (SEM). Chip mapping and the imaging of serial sections was facilitated by SmartSEM (Zeiss) and Atlas 5 (FIBICS) software packages. Maps of all the serial sections on the silicon chips were collected at 500nm/px. Laser marks could be identified in the resin boundary at low magnification, demarcating the ROI in each section. The ROI was identified and captured across 32 serial section scanning electron micrographs. ROIs were imaged at 2nm/px using an electron backscatter detector (Gatan). In high-vacuum mode, a 3keV beam in high-current mode with a 30 μm aperture at a working distance of 9 mm was found to produce signal from which we could resolve membrane and organelle (e.g., microtubule) ultrastructure. Sections were aligned using Photoshop (Adobe) and elastic alignment functions embedded in Fiji (NIH)^8^. The orthogonal projection (e.g., x-z) of the SEM data was used to correlate the fluorescence data and produce overlay using Imaris.

### Cellular cryo-electron tomography (cryo-ET) data acquisition and tomogram reconstruction

*Tfam^+/−^* MEFs (TFAM Het MEF) and SV40 mouse embryonic fibroblasts (SV40 MEF) were cultured on R ¼ Carbon 200-mesh gold electron microscopy grids (Quantifoil Micro Tools) and plunge frozen in a liquid ethane/propane mixture using a Vitrobot Mark 4 (Thermo Fischer Scientific). Thin (120-200 nm) vitrified lamellae were prepared by Focused Ion Beam milling using an Aquilos dual-beam FIB/SEM instrument (Thermo Fisher Scientific) following a modified version of a previous manual cryo-preparation workflow^9^. Grids containing lamellae were transferred into a 300keV Titan Krios microscope (Thermo Fisher Scientific) with a K2 Summit direct electron detector camera (Gatan). Low magnification tilt series were acquired using SerialEM software^10^ with 2° steps between −60° and +60°. Individual tilt were collected with a pixel size of 7.151 Å and a defocus range of −25.9 to −28.1. The dose per tilt was 0.189 e/Å^2^, and the total accumulated dose for the tilt series was under 11.5 e/Å^2^. Regions of multivesicular bodies were selected for high magnification data collection. High magnification tilt series were acquired using SerialEM software with 2° steps between −60° and +60°. Individual tilts were collected with a pixel size of 3.106 Å and a defocus range of −15.0 to −17.2. The total dose per tilt was 0.905 e/Å^2^, and the total accumulated dose for the tilt series was under 55 e/Å^2^. Alignment of tilt series were performed in IMOD^11^ using patch tracking. Cryotomograms were reconstructed using weighted back projection in IMOD.

### Quantification of multivesicular bodies in tomograms

High magnification tomograms were used for quantification of multivesicular bodies (MVBs). MVBs were identified as non-mitochondrial multi-membrane structures in tomograms and were counted only if >1/2 of the MVB was visible in the field of view of the tomogram to avoid overlapping picks. The total number of MVBs was counted per individual tomogram in each condition. Total tomograms for WT SV40 MEF n= 15 and TFAM Het n= 27. Mean values for MVB in WT SV40 MEF = 0.66 and TFAM Het = 1.03.

### Quantitative PCR

To quantify messenger RNA transcript abundance, RNA was extracted from cells in 6 cm dishes using the RNeasy kit (Qiagen # 74106) according to manufacturer instructions. The optional steps were included during RNA extraction, and elution time during was extended to 10 minutes to improve yield. Reverse transcription was achieved using High Capacity cDNA Kit (Thermo Fisher # 4368814) with 2 μg RNA input into each reaction. Equal amounts of complementary DNA and the indicated primers (Extended Data Table 1, purchased from Eaton Biosciences) were used for quantitative PCR (qPCR) using Fast SYBR Green Master Mix (Thermo Fisher #4364346). For each biological sample, three technical replicates were performed and normalized against the *Beta actin* or *Gapdh* threshhold cycle (Ct) value to calculate ΔCt. The ΔCt of each sample was then compared to the ΔCt of the control sample to generate the ΔΔCt value. Relative expression was then analyzed using the 2-ΔΔCt method and the relative fold change was plotted with the control samples given a value of 1.0. To quantify mtDNA, cells were suspended in 50 mM NaOH and boiled for 1 hour by neutralization by 1/10 volume of 1 M Tris-HCl (pH 8.0). DNA samples were diluted to 10 ng μl^-1^ and subjected to qPCR analysis using D-loop (two D-loop and ND4 primers, Supplementary Table 1) and 18S primers (Supplementary Table. 1) to amply mtDNA and nDNA, respectively. Three technical replicates were performed for each biological sample and normalized against nuclear 18S value. Relative copy number was analyzed using a 2-ΔΔCt method and the control mtDNA abundance was given a value of 100%.

### Ratiometric Flow Cytometry to Measure Mitophagy Flux

U2OS cells with stable expression of a mCherry-GFP-FIS1 tandem construct (a generous gift from Dr. Ian Ganley’s group^12^) were used to measure mitophagy via ratiometric flow cytometry as has been described previously^13^. Cells were transfected or treated with different mitophagy inducing drugs 24 hours prior to flow cytometry analysis on an Aria Fusion Cell Sorter using lasers set at 488nM and 561nM. Debris, dead cells, and doublets were omitted via side/forward scatter profiling. A “Tandem+” gate was set using unstained cells with the gate set to 1% “tandem+”. FlowJo software was then used to analyze the mCherry/GFP ratio within the “tandem+” gate. A “mitophagy” gate was set based on the mCherry/GFP ratio of the Bafilomycin-A1 treated sample set to 5%.

### Image analysis, data quantification, and statistics

Imaging data was quantified using Fiji (ImageJ). All statistical analyses and graphs were generated using GraphPad Prism 8 software. Differences >0.05 by unpaired, two tailed student’s t test were considered significant. *P < 0.05; **P < 0.01; ***P < 0.001; ****P < 0.0001; NS, not significant (P > 0.05).

### Quantification of cytoplasmic nucleoids and overlap with cGAS/RAB7

Cells were selected for imaging if they displayed puncta positive for cGAS and DNA or mtDNA FISH. The mitochondria channel was masked using automatic thresholding, and this mask was subtracted from the other channels using the image calculator. The mtDNA FISH and other channels were then thresh-holded and instances of overlap were scored (1 for overlap, 0 for no overlap).

### Quantification of nucleoid size and EdU intensity

Nuclear signal was first subtracted from the DNA channel. The remaining DNA signal was masked using automatic thresholding, and nucleoids were then measured using “Analyze particles.” Nucleoids were considered enlarged if larger than 0.4 μm^2^. To quantify EdU intensity, EdU fluorescence intensity was measured within the DNA mask. To quantify EdU/DNA ratio, the EdU intensity within the DNA mask was divided by the DNA intensity and plotted as a function of nucleoid size.

### Quantification of mitochondrial and ER-associated actin

Nucleoid size was first measured as described above, and nucleoids larger than 0.3 μm^2^ were classified as enlarged. For each cell (N=5), line scans (1.5 μm) were then drawn through three normal and three enlarged nucleoids. The peak fluorescence intensity of each actin probe was measured and normalized to the mitochondrial or ER fluorescence intensity within 164 nm of the actin peak.

## Acknowledgements

The authors gratefully acknowledge Drs. Jodi Nunnari, Susan Kaech, Phillip West, and Zheng Wu for their impactful ideas and suggestions. We also thank Dr. Henry Higgs (Dartmouth) for the generous gift of INF2 knockout cells and Drs. Rebecca Gilson and Cayla Miller (Salk Biophotonics Core) for help aligning the CLEM datasets. This work was supported by NIH R01 AR069876 to G.S.S., who also holds the Audrey Geisel Chair in Biomedical Science, and the Allen-AHA Initiative in Brain Health and Cognitive Impairment Award 19PABH134610000H, NIH 1K99GM141482 and George E. Hewitt Foundation for Medical Research Postdoctoral Fellowship to L.E.N., Paul F. Glenn Foundation for Medical Research Postdoctoral Fellowship to N.T., and NIH 1F32GM137580 to C.R.S. This work was supported by the Waitt Advanced Biophotonics Core Facility of the Salk Institute with funding from NIH-NCI CCSG: P30 014195 and the Waitt Foundation, as well as the Yale University School of Medicine Center for Cellular and Molecular Imaging. This work was supported by 5R00CA245187 and 5R00CA245187-04S1 as well as the Flow Cytometry Core Facility of the Salk Institute with funding from NIH-NCI CCSG: P30 014195 and Shared Instrumentation Grant S10-OD023689 (Aria Fusion cell sorter).

The models in Fig. 4G and Extended Data Fig. 8 were created using Biorender.com.

## Author Contributions

L.E.N., N.T., U.M., and G.S.S. planned the experimental design and data analysis; L.E.N., N.T., C.R.S., G.R.R., J.A.C., and I.L. performed the experiments; L.E.N., N.T., C.R.S., and G.R.R. performed data analysis and quantification; L.E.N. and S.W.N. performed the CLEM experiments and analyzed the data, M.M. and D.A.G. performed the cryo-EM experiments, analyzed the data, and composed the figures for those experiments, S.R. and C.G.T. performed the mitophagy experiments, analyzed the data, and composed the figures for those experiments, U.M. and G.S.S. supervised the study; L.E.N. composed the figures and videos, and L.E.N., U.M., and G.S.S. wrote the manuscript with input from the rest of the authors.

## Data Availability Statement

All data are provided with the paper, and any raw data or reagents will be provided by the corresponding authors upon reasonable request.

## Additional Information

**Extended Data Table 1.**
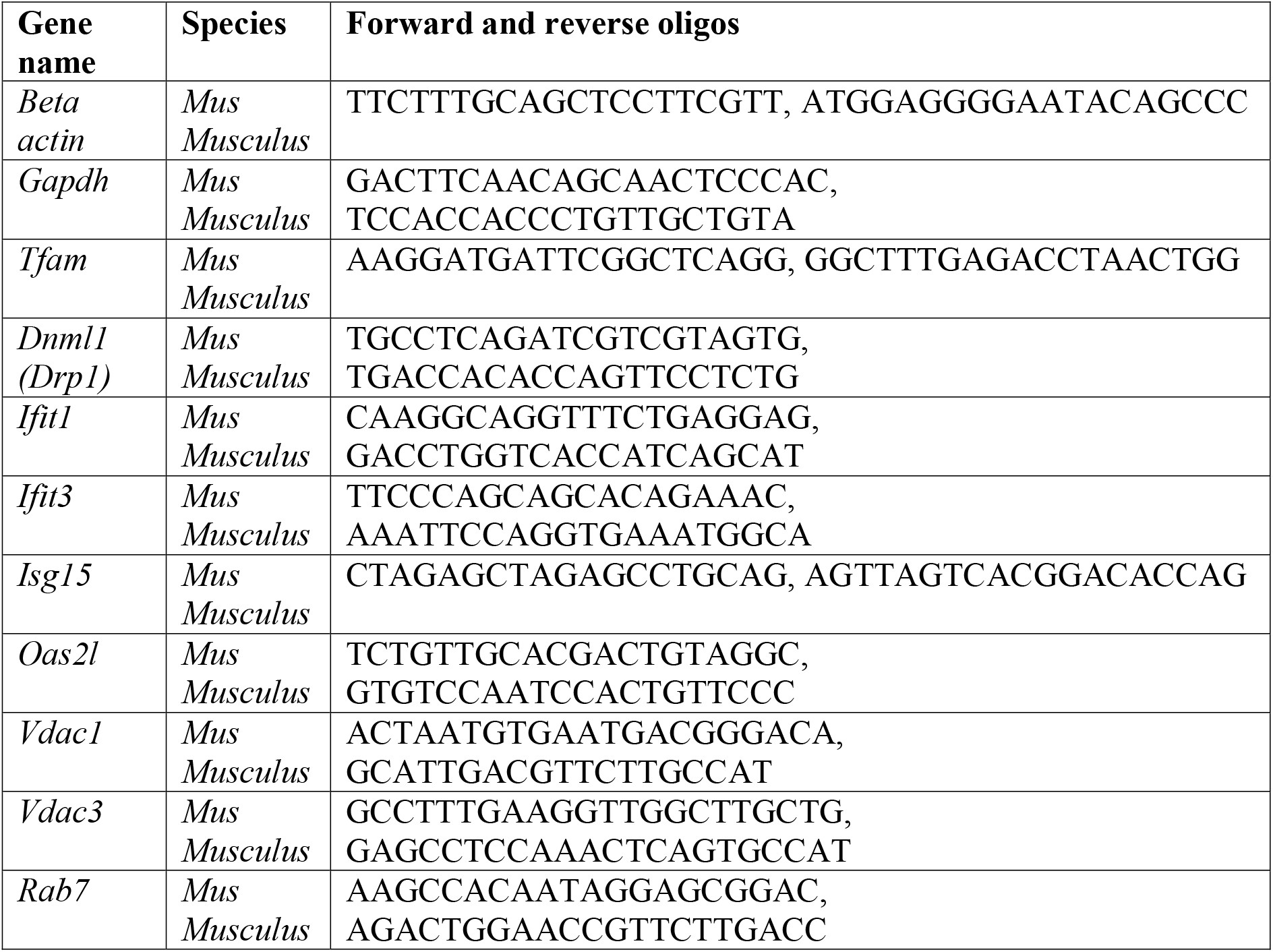
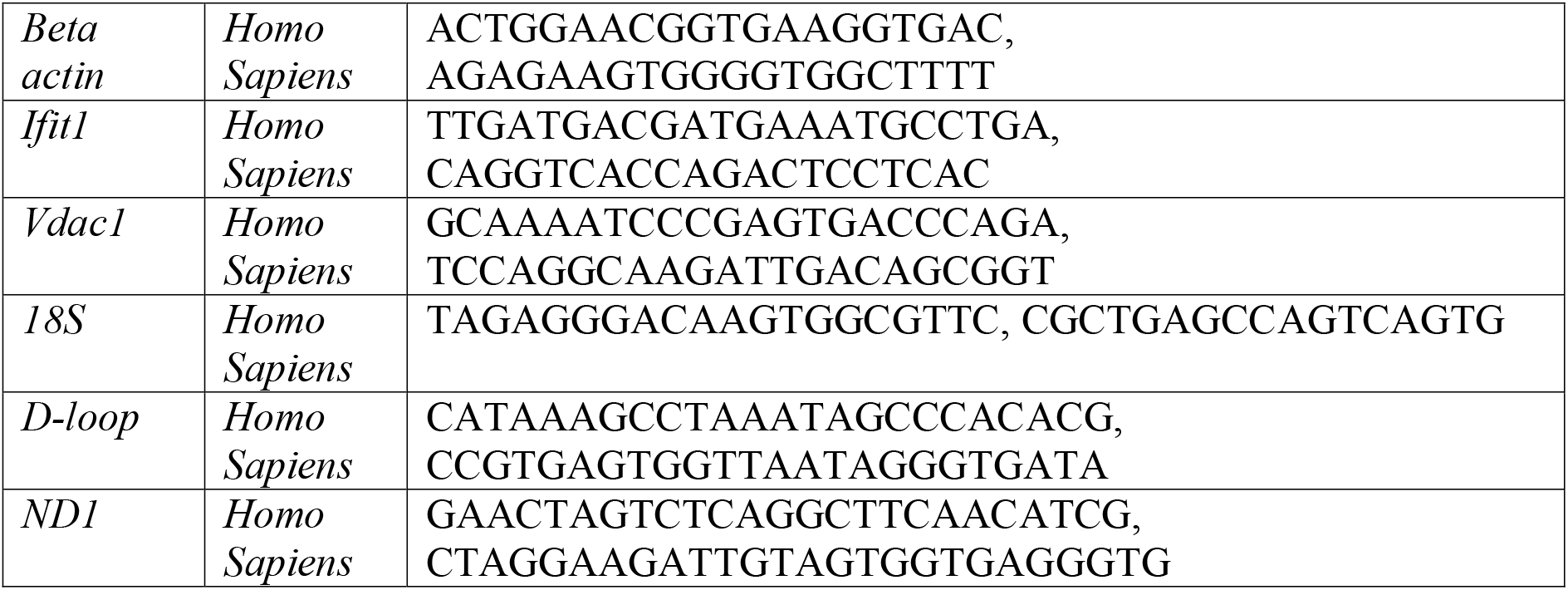
PCR primer sequences.

**Extended Data Table 2.**
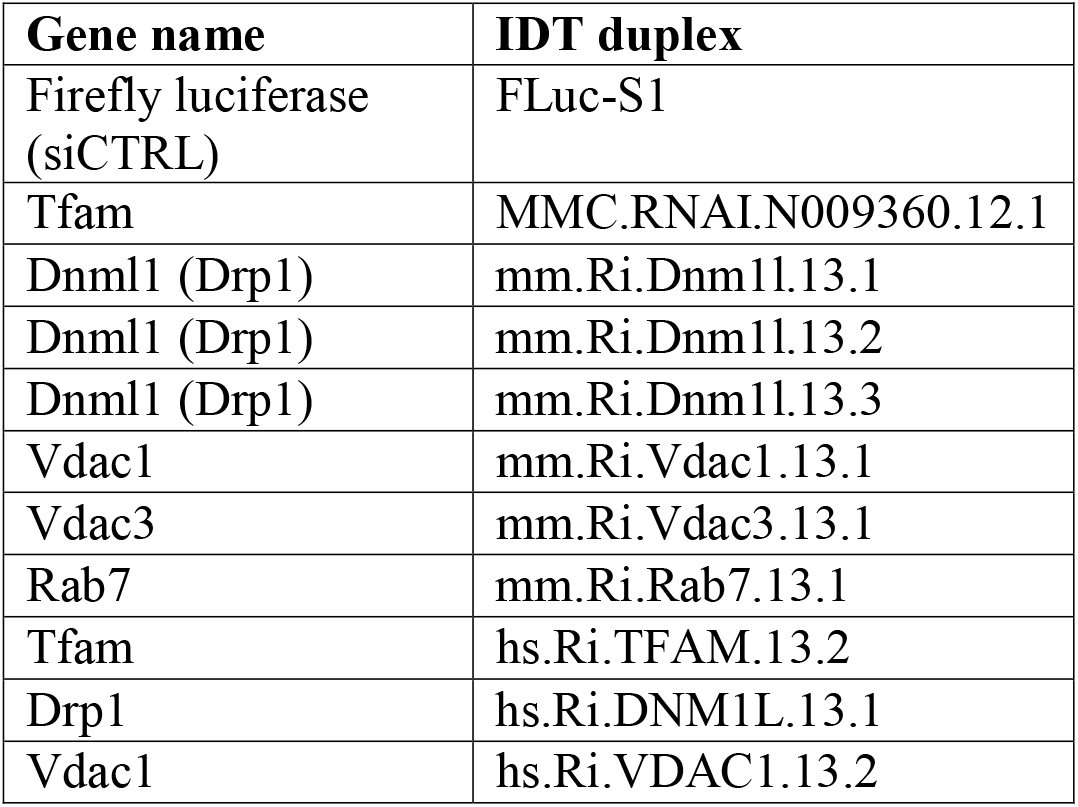
List of siRNAs.

| **Gene name** | **IDT duplex** |
| --- | --- |
| Firefly luciferase (siCTRL) | FLuc-S1 |
| Tfam | MMC.RNAI.N009360.12.1 |
| Dnm1l (Drp1) | mm.Ri.Dnm1l.13.1 |
| Dnm1l (Drp1) | mm.Ri.Dnm1l.13.2 |
| Dnm1l (Drp1) | mm.Ri.Dnm1l.13.3 |
| Vdac1 | mm.Ri.Vdac1.13.1 |
| Vdac3 | mm.Ri.Vdac3.13.1 |
| Rab7 | mm.Ri.Rab7.13.1 |
| Tfam | hs.Ri.TFAM.13.2 |
| Drp1 | hs.Ri.DNM1L.13.1 |
| Vdac1 | hs.Ri.VDAC1.13.2 |

## Extended Data Figure Legends

**Extended Data Figure 1.**
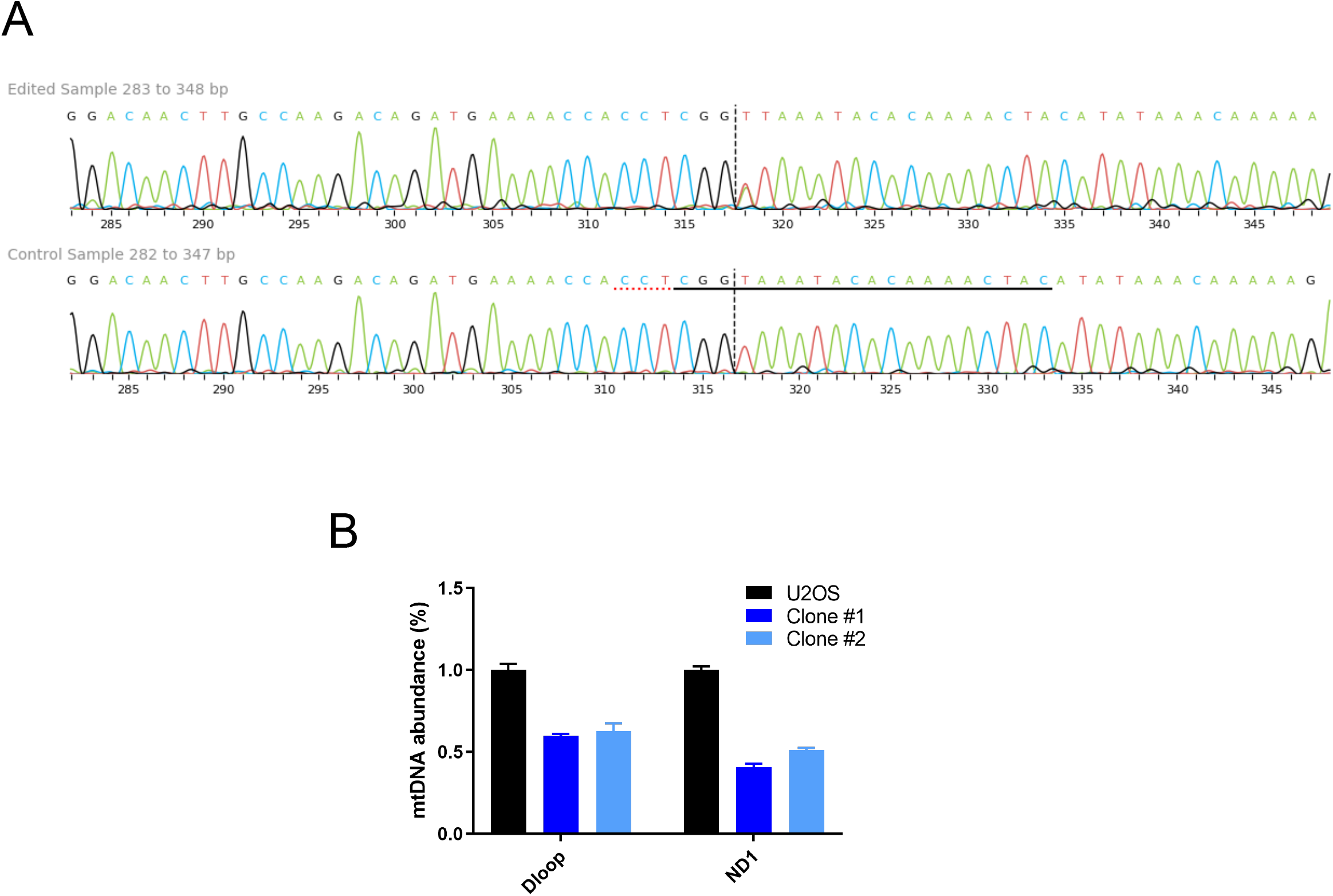
Characterization of TFAM-deficient U2OS cells derived using CRISPR. **A)** Sequencing of U2OS TFAM-deficient clones, confirming insertion and frameshifting in exon 1. **B)** mtDNA abundance (relative mtDNA copy number) by qPCR with D-loop and ND1 primers, normalized to nuclear 18S. Results were reproducible across three independent experiments.

**Extended Data Figure 2.**
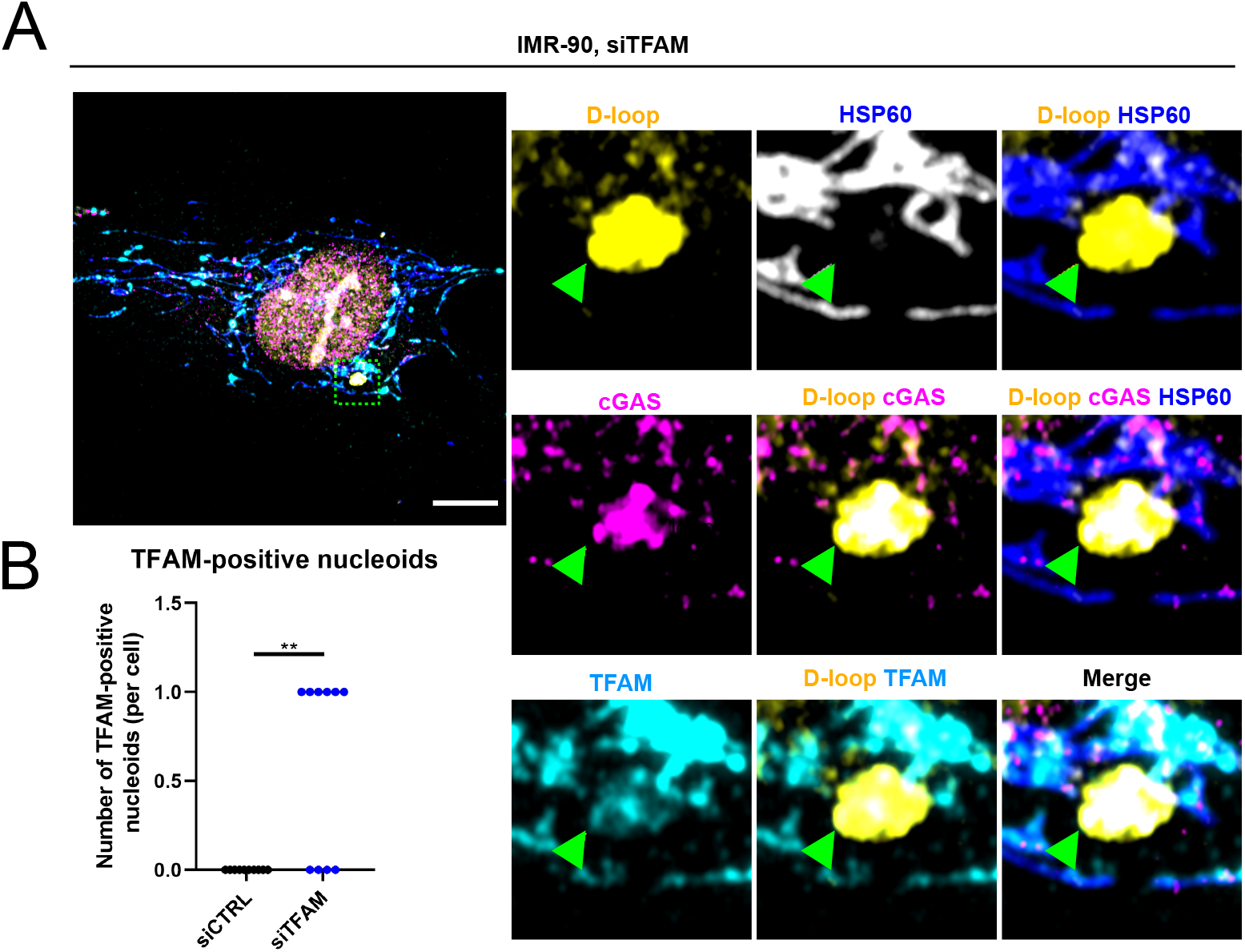
TFAM-bound nucleoids are released in TFAM knock-down cells. **A)** Airyscan imaging of mtDNA FISH using a D-loop probe, followed by immunofluorescence against cGAS, HSP60, and TFAM, in IMR-90 cells transfected TFAM siRNA. **B)** Quantification of cytoplasmic nucleoids positive for TFAM as in Figure 1D. N=10 for both conditions.

**Extended Data Figure 3.**
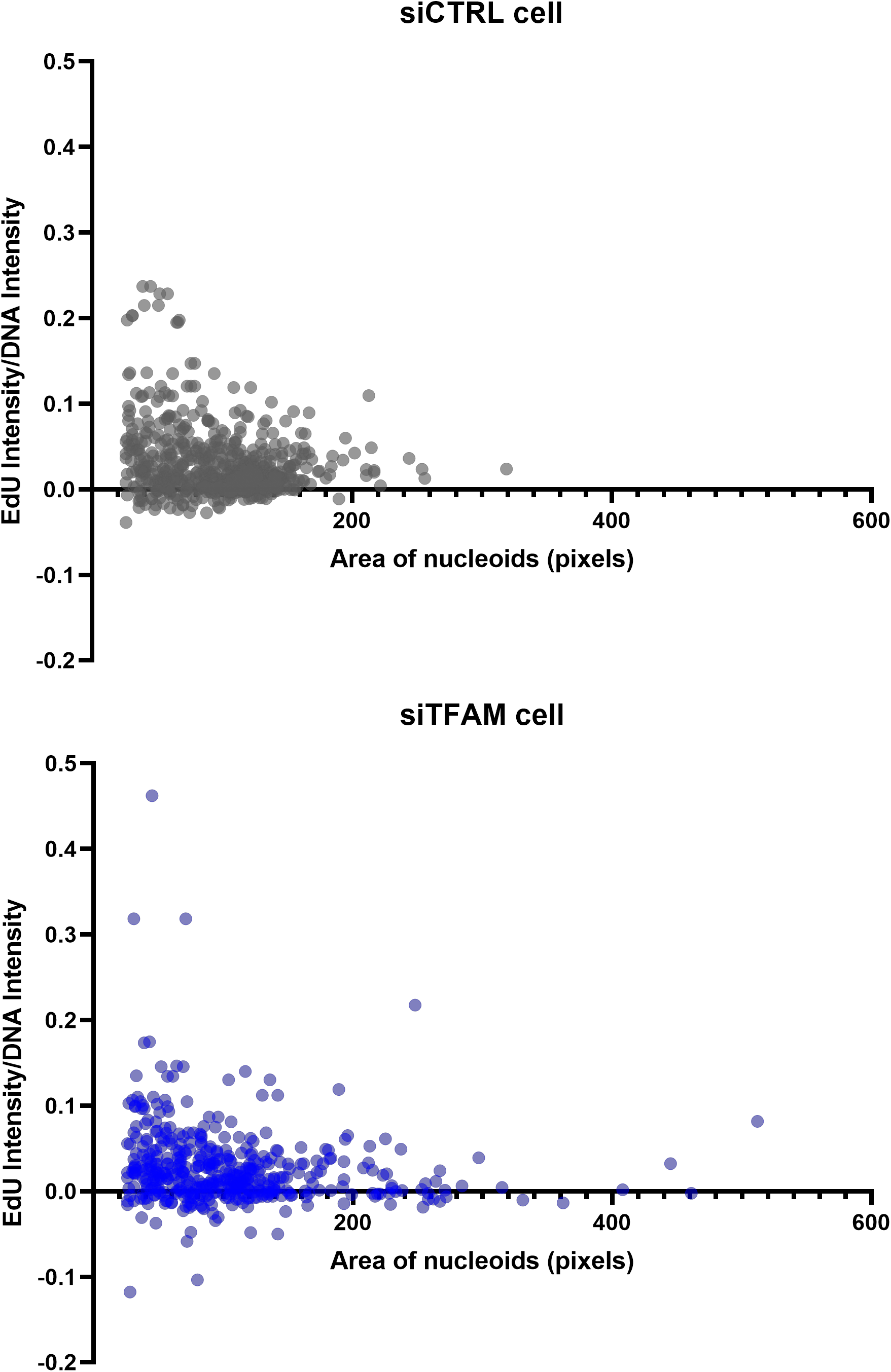
EdU incorporation is inversely correlated with nucleoid size in TFAM-deficient cells. EdU intensity within nucleoids was quantified in siCTRL and siTFAM MEFs as in Figure 2B and normalized to DNA fluorescence intensity. The ratio of EdU to DNA fluorescence intensity is plotted against nucleoid size, indicating that EdU intensity diminishes as nucleoid size increases. Data from one control cell and one TFAM-depleted cell are shown.

**Extended Data Figure 4.**
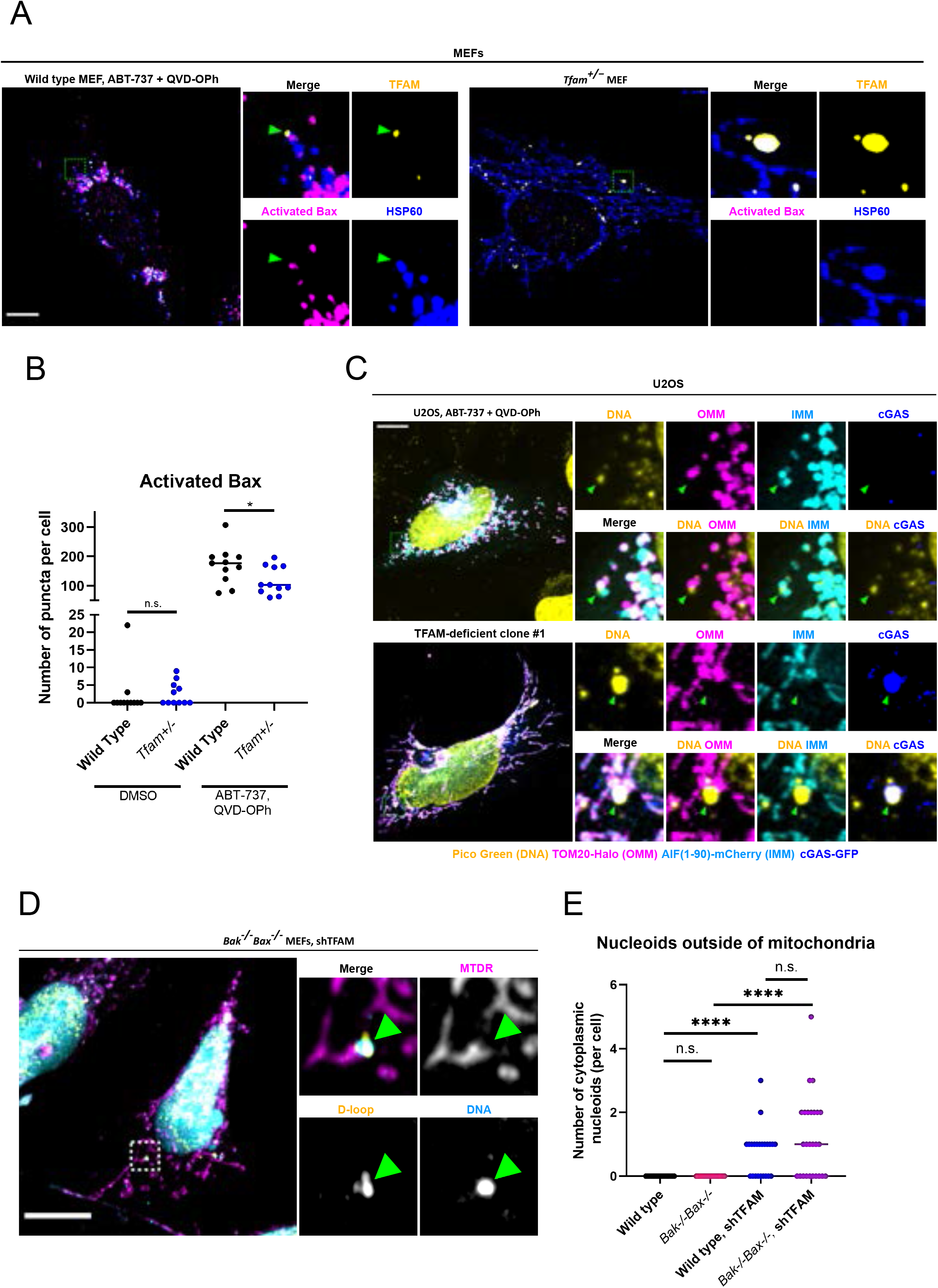
Enlarged nucleoids are not released through BAX pores. **A)** Airyscan imaging of immunofluorescence against activated BAX alongside TFAM and HSP60 in primary wild type and *Tfam^+/−^* MEFs. As a positive control, BAX-mediated mtDNA release was induced by treatment with ABT-737 (10 μm) plus QVD-OPh (20 μM) for 4 hours. **B)** Quantification of **(A).** The number of BAX puncta were counted per cell. N=11 cells for each condition. **C)** Airyscan imaging of TFAM-deficient U2OS cells expressing AIF(1-90)-mCherry (IMM), TOM20-Halo (OMM), and cGAS-GFP, along with immunofluorescence against DNA. ABT-737 and QVD-OPh treatment was used as a positive control for IMM herniation beyond the OMM. Results were reproducible across three independent experiments. **D)** Airyscan imaging of mtDNA using a D-loop probe, followed by immunofluorescence against TOM20 in *Bak^−/−^Bax^−/−^* MEFs stably expressing TFAM shRNA. **E)** Quantification of cytoplasmic nucleoids per cell. N=21 for both wild type and *Bak^−/−^Bax^−/−^* MEFs, N=23 for WT shTFAM, N=26 for *Bak^−/−^Bax^−/−^* shTFAM.

**Extended Data Figure 5.**
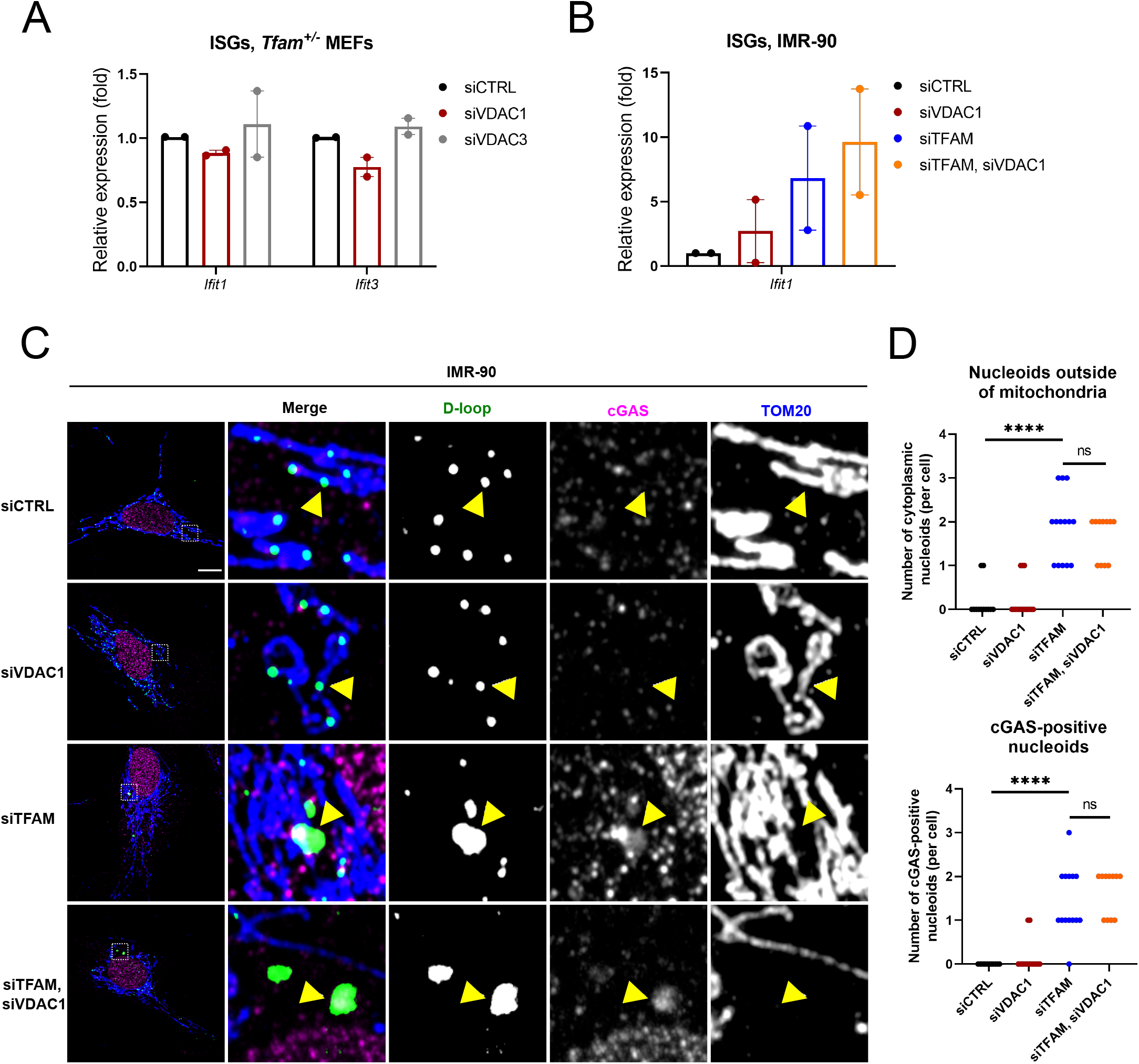
Enlarged nucleoids are not released through VDAC pores. **A)** qRT-PCR of primary *Tfam^+/−^MEFs* transfected with control, VDAC1, or VDAC3 siRNAs, normalized to beta actin. Dots represent biological replicates (N=2). **B)** qRT-PCR of IMR-90 cells transfected with control, TFAM, VDAC1, or TFAM and VDAC1 siRNAs, normalized to beta actin. Dots represent technical replicates (N=2). **C)** Airyscan imaging of mtDNA FISH using a D-loop probe, followed by immunofluorescence against cGAS and TOM20, in IMR-90 cells transfected as in B. **D)** Quantification of released and cGAS-positive nucleoids as described in Fig. 1D. N=12 for siCTRL, N=15 for siVDAC1, N=14 for siTFAM, N=11 for siTFAM, siVDAC1.

**Extended Data Figure 6.**
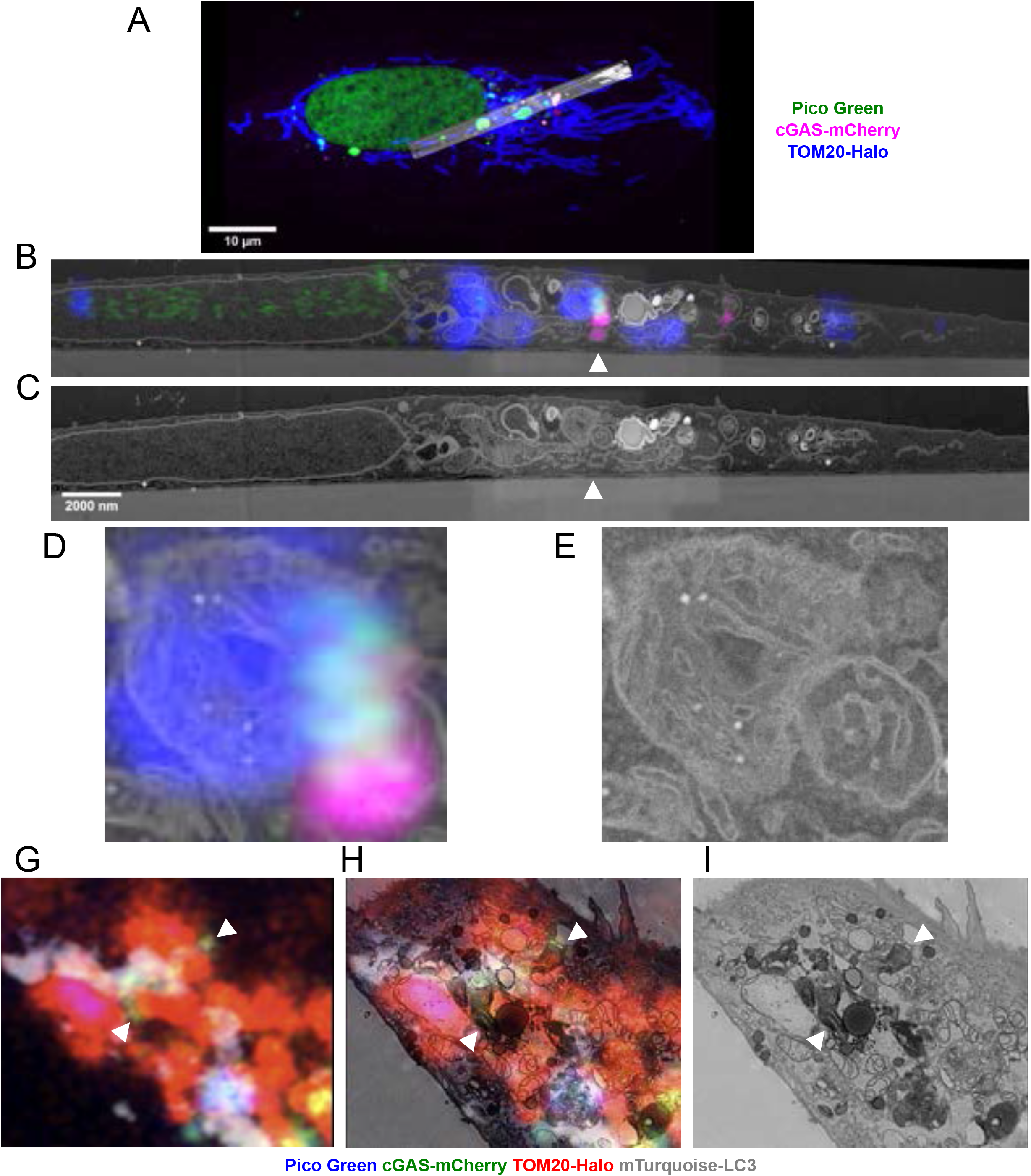
Enlarged nucleoids are released from mitochondria in a membrane-bound compartment, and cGAS overlaps with membrane-bound compartments in TFAM-deficient cells. TFAM-deficient U2OS cells (clone #1) were imaged live using airyscan microscopy until a cGAS-positive nucleoid was observed to be exiting mitochondria. The cell was fixed on the microscope stage and processed for correlative light and electron microscopy (see methods). **A)** Overview of cell showing EM data aligned to fluorescence data. **B)** One EM slice aligned to fluorescence data is shown, arrowhead indicates mtDNA being released. **C)** Same as B showing only EM data. **D)** Blowup of ROI from aligned data. **E)** Blowup of aligned data from EM. **F)** Fluorescence data from a second cell fixed on the stage. **G)** One EM slice aligned to the fluorescence data is shown. **H)** One EM slice is shown. Arrowheads indicate cGAS-mCherry fluorescence signal correlated with membrane-bound compartments. The mtDNA release event for this cell was not captured on the EM sections obtained from this cell due to difficulties with serial sectioning.

**Extended Data Figure 7.**
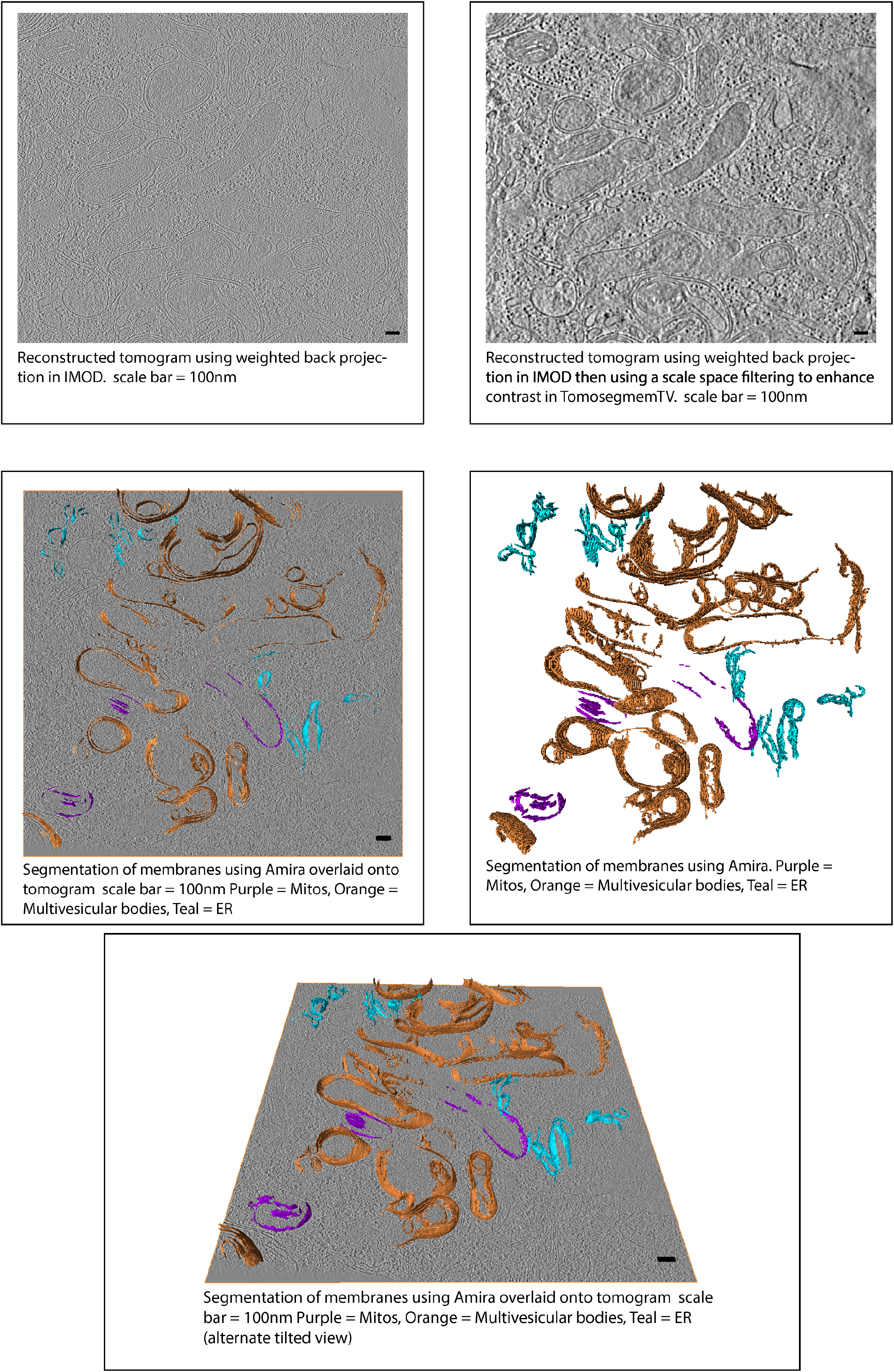
Segmentation of organelles in cryo-EM tomograms of *Tfam^+/−^* MEFs. A representative tomogram slice of a *Tfam^+/−^* MEF cell imaged by cellular cryo-electron tomography. Membranes were segmented and colored orange for multivesicular bodies, purple for mitochondria, and teal for ER. Scale bar = 100 nm.

**Extended Data Figure 8:**
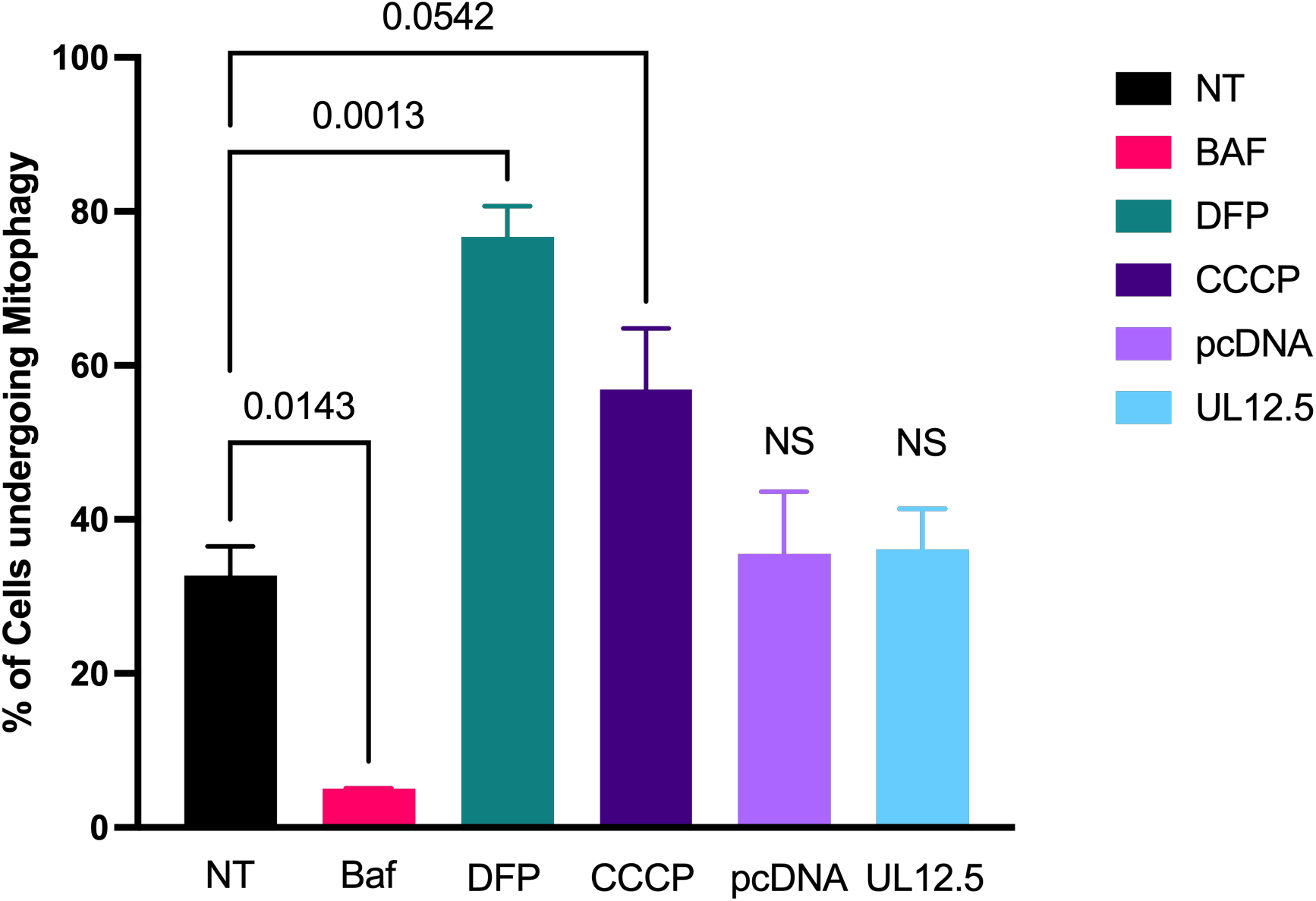
No change in mitophagy flux with expression of HSV-1 UL12.5. Quantification of mitophagy via ratiometric flow cytometry performed in U2OS cells with stable expression of mCherry-GFP-Fis1. Cells were either transfected with pcDNA3.1, UL12.5, or treated with Bafilomycin-A1 (20nM), Deferiprone (DFP 1 mM), or CCCP (20μM) 24 hours before flow cytometry analysis. The data are shown as the average ratio of mCherry/GFP ± SEM from biological replicate experiments (N of 2-3).

**Extended Data Figure 9.**
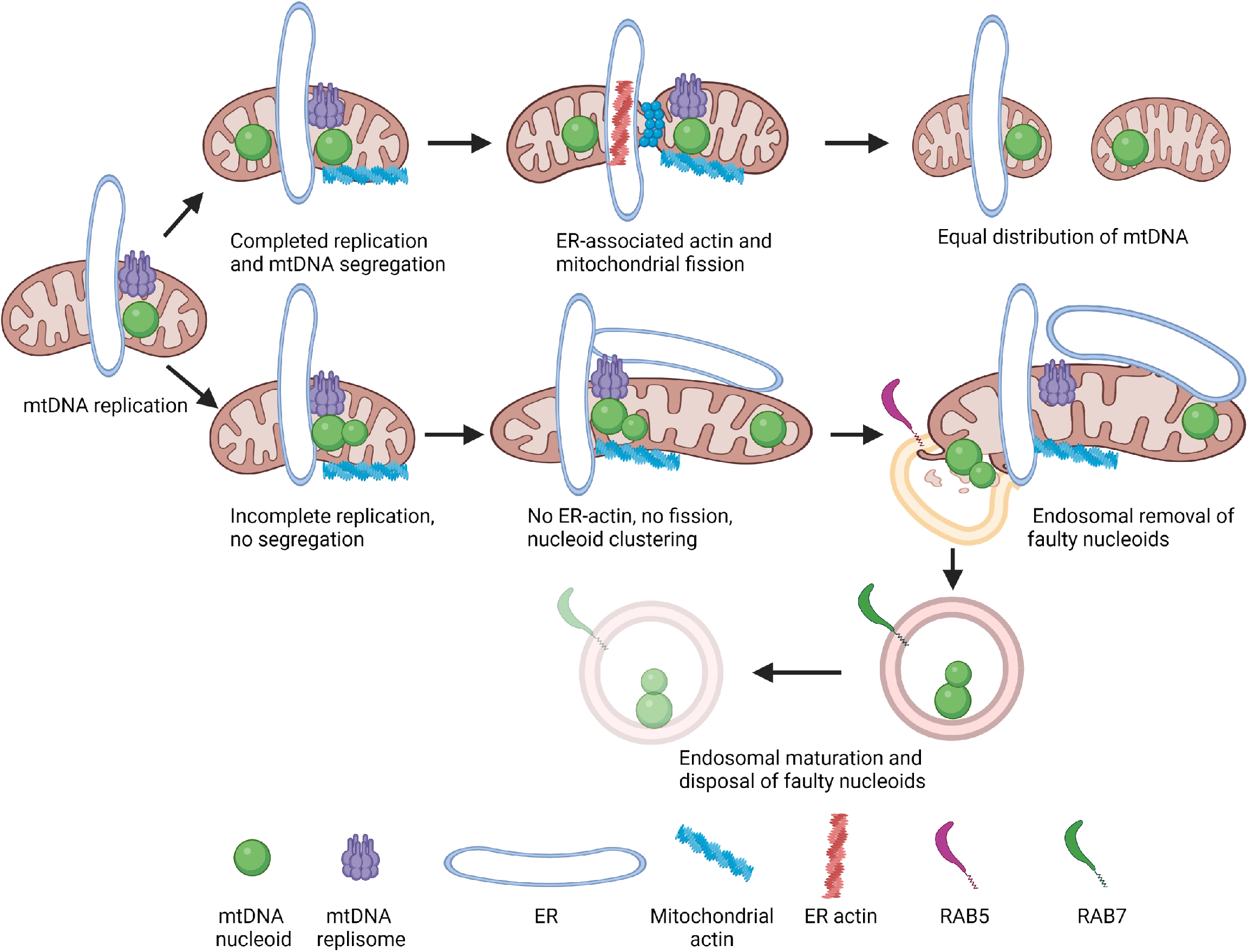
Model summarizing endosome-mediated removal and disposal of dysfunctional nucleoids. Following mtDNA replication, a signal is sent from mitochondria to the ER allowing for polymerization of ER-associated actin, which allows newly replicated nucleoids to segregate via mitochondrial fission. If problems arise during mtDNA replication or termination, no signal is sent to the ER, and actin does not associate with the ER. If nucleoids are unable to properly segregate through mitochondrial fission, a fission checkpoint is enacted (to wait for completion of mtDNA segregation), and clustered nucleoids accumulate at sites of replication. If not rectified, the dysfunctional nucleoids are extracted via endosomes. This process activates cGAS/STING signaling initially, but the proinflammatory nucleoids are eventually disposed of by late endosomes.

**Supplemental Video 1. Live cell imaging of mtDNA release in *Tfam^+/−^*AMEFs.** Airyscan live cell imaging of Pico Green (DNA) and TOM20-mCherry (OMM) in a live, immortalized *Tfam^+/−^* MEF cell. Arrow indicates release. Scale bar = 5 μm. This video corresponds to Fig. 1A.

**Supplemental Video 2. Live cell imaging of mtDNA release and colocalization with cGAS-mCherry.** Airyscan live cell imaging of Pico Green (DNA), cGAS-mCherry, and TOM20-Halo (OMM) in a live TFAM-deficient U2OS cell. Arrow indicates mtDNA release. Scale bar = 10 μm. This video corresponds to Fig. 1B.

